# SARS-CoV-2 infection induces germinal center responses with robust stimulation of CD4 T follicular helper cells in rhesus macaques

**DOI:** 10.1101/2020.07.07.191007

**Authors:** Sonny R. Elizaldi, Yashavanth Shaan Lakshmanappa, Jamin W. Roh, Brian A. Schmidt, Timothy D. Carroll, Kourtney D. Weaver, Justin C. Smith, Jesse D. Deere, Joseph Dutra, Mars Stone, Rebecca Lee Sammak, Katherine J. Olstad, J. Rachel Reader, Zhong-Min Ma, Nancy K. Nguyen, Jennifer Watanabe, Jodie Usachaenko, Ramya Immareddy, JoAnn L. Yee, Daniela Weiskopf, Alessandro Sette, Dennis Hartigan-O’Connor, Stephen J. McSorley, John H. Morrison, Nam K. Tran, Graham Simmons, Michael P Busch, Pamela A. Kozlowski, Koen K.A. Van Rompay, Christopher J. Miller, Smita S. Iyer

## Abstract

CD4 T follicular helper (T_fh_) cells are important for the generation of long-lasting and specific humoral protection against viral infections. The degree to which SARS-CoV-2 infection generates T_fh_ cells and stimulates the germinal center response is an important question as we investigate vaccine options for the current pandemic. Here we report that, following infection with SARS-CoV-2, adult rhesus macaques exhibited transient accumulation of activated, proliferating T_fh_ cells in their peripheral blood on a transitory basis. The CD4 helper cell responses were skewed predominantly toward a T_h_1 response in blood, lung, and lymph nodes, reflective of the interferon-rich cytokine environment following infection. We also observed the generation of germinal center T_fh_ cells specific for the SARS-CoV-2 spike (S) and nucleocapsid (N) proteins, and a corresponding early appearance of antiviral serum IgG antibodies but delayed or absent IgA antibodies. Our data suggest that a vaccine promoting Th1-type Tfh responses that target the S protein may lead to protective immunity.

## INTRODUCTION

As of July 6^th^, 2020, SARS-CoV-2 has resulted in more than 11.6 million infections and more than half a million deaths, globally (1, 2). Unanticipated post-infection complications, such as multisystem inflammatory syndrome pose a serious threat (3). An effective vaccine is paramount, and there are several SARS-CoV-2 vaccine candidates, including vaccines based on platform technologies that have shown promise against the coronaviruses that cause SARS and MERS, in various phases of human testing worldwide (4-6). The most effective vaccines induce antibodies that provide long-term protection, exhibit specificity and avidity for the antigen or subunit of the antigen, and are capable stopping replication or otherwise inactivating the pathogen (7). Vaccines using attenuated virus elicit the most persistent antibody responses; therefore, understanding the immunological mechanisms characteristic of SARS-CoV-2, specifically immune responses associated with production of antibodies against the spike glycoprotein, is foundational to the selection of a vaccine capable of abating the pandemic (8, 9).

Generation of persistent immunity hinges on CD4 T follicular helper cells (T_fh_). We and others have demonstrated that peripheral CD4 T_fh_ cells predict antibody durability in the context of HIV and influenza vaccines (10-12). The impact of SARS-CoV-2 infection on the generation of T_fh_ cells is currently unknown. This is a detrimental gap in knowledge as understanding early correlates of durable antibodies, specifically those that circulate in peripheral blood, will aid in the ultimate selection of effective vaccine candidates. SARS-CoV-2-specific CD4 T cells responding to spike proteins have been observed in the peripheral blood samples of recovered patients (13, 14). Similar observations have been made with the 2002 SARS-CoV virus (15, 16), and studies in mouse models have demonstrated a critical role for CD4 T cells in viral clearance (6). Together, these data emphasize the need to understand CD4 T_fh_ responses following SARS-CoV-2 infection. While several recent studies have reported on T cell dynamics in peripheral blood of patients (17-21), early immune responses, particularly in lymphoid and respiratory tissues, are challenging to study in humans. Rhesus macaques have emerged as a robust model for SARS-CoV-2 (22-27). Because healthy rhesus macaques infected with SARS-CoV-2 resist immediate re-challenge with the virus (24, 27), we hypothesized that understanding the CD4 T_fh_ and germinal center (GC) response following exposure to SARS-CoV-2 will provide a framework for understanding immune mechanisms of protection thereby providing evidence-based data on which to select an effective vaccine.

Here we report that SARS-CoV-2 infection triggered acute shifts in peripheral innate myeloid cells in adult rhesus macaques. Notably, on Day 2 post viral exposure we observed a dramatic rise in pro-inflammatory monocytes and decline in plasmacytoid dendritic cells (pDCs) in peripheral blood. This change was only transient and began to subside on Day 4 in conjunction with rapid resolution of systemic inflammation early during the course of infection, consistent with mild clinical symptoms. Perhaps more pertinent to SARS-CoV-2 as a respiratory virus, infection elicited robust GCs with SARS-CoV-2-reactive T_fh_ cells within the mediastinal lymph nodes. Additionally, CD4 T_fh_ cells - specifically T_h_1-T_fh_ - were observed in peripheral blood following infection. The data suggest that vaccine platforms inducing T_h_1-T_fh_ responses are likely to succeed in eliciting durable humoral responses. Our findings only begin to bridge the gap in knowledge that exists in understanding the immune response triggered by SARS-CoV-2 - specifically T_fh_ and GC responses - and further investigation will provide a solid framework for rational vaccine design and selection.

## RESULTS

### Experimental Design

To achieve our primary objective of assessing whether SARS-CoV-2 elicits T_fh_ cells and germinal center responses, we challenged eight adult rhesus macaques (four to five years of age, additional details provided in (**Table S1**) with a high-dose of SARS-CoV-2 (2×10^6^ PFU; corresponding to 2×10^9^ vRNA). Virus was administered via the intranasal, intratracheal, and ocular routes. Infection was determined by monitoring viral RNA (vRNA) in nasal washes using qRT-PCR. Of the eight animals: four animals were infected with SARS-CoV-2 and did not receive any human plasma serving as an infected-only group (Infected), two animals were infected and infused with COVID-19 convalescent human plasma (I+CP), 24 hours following inoculation, and two animals were infected and infused with an identical volume of normal plasma (I+NP), with no antibodies against SARS-CoV-2. (**Figure 1A**). Pooled CP demonstrated a neutralizing titer of 1:1,149; but due to the expected ∼50-fold dilution after infusion (estimated based on infusing 4 ml/kg and ∼ 20% extracellular fluid), neutralizing activity in macaque sera 24 hours post infusion fell below the limit of detection (1:40) of the neutralization assay. Consistently, CP administration did not blunt acute viral replication kinetics and high levels of vRNA were observed in all animals (**Figure 1B**). Histopathological lesions of the lungs between day 11 - day 14 post infection confirmed multifocal to locally extensive interstitial pneumonia of mild to moderate severity in all infected animals (**Figure S1A**). However, these histological changes were not accompanied by fever, weight loss, or any other signs of clinical disease (**Figure S1B**). One animal developed dermatitis but only a few animals were found sneezing (**Table S1**). None of the animals developed acute respiratory distress. In summary, infection of healthy adult rhesus macaques with SARS-CoV-2 resulted in high viremia but generally produced no overt signs of clinical illness, providing a framework to investigate development of protective immune responses.

**Figure 1.**
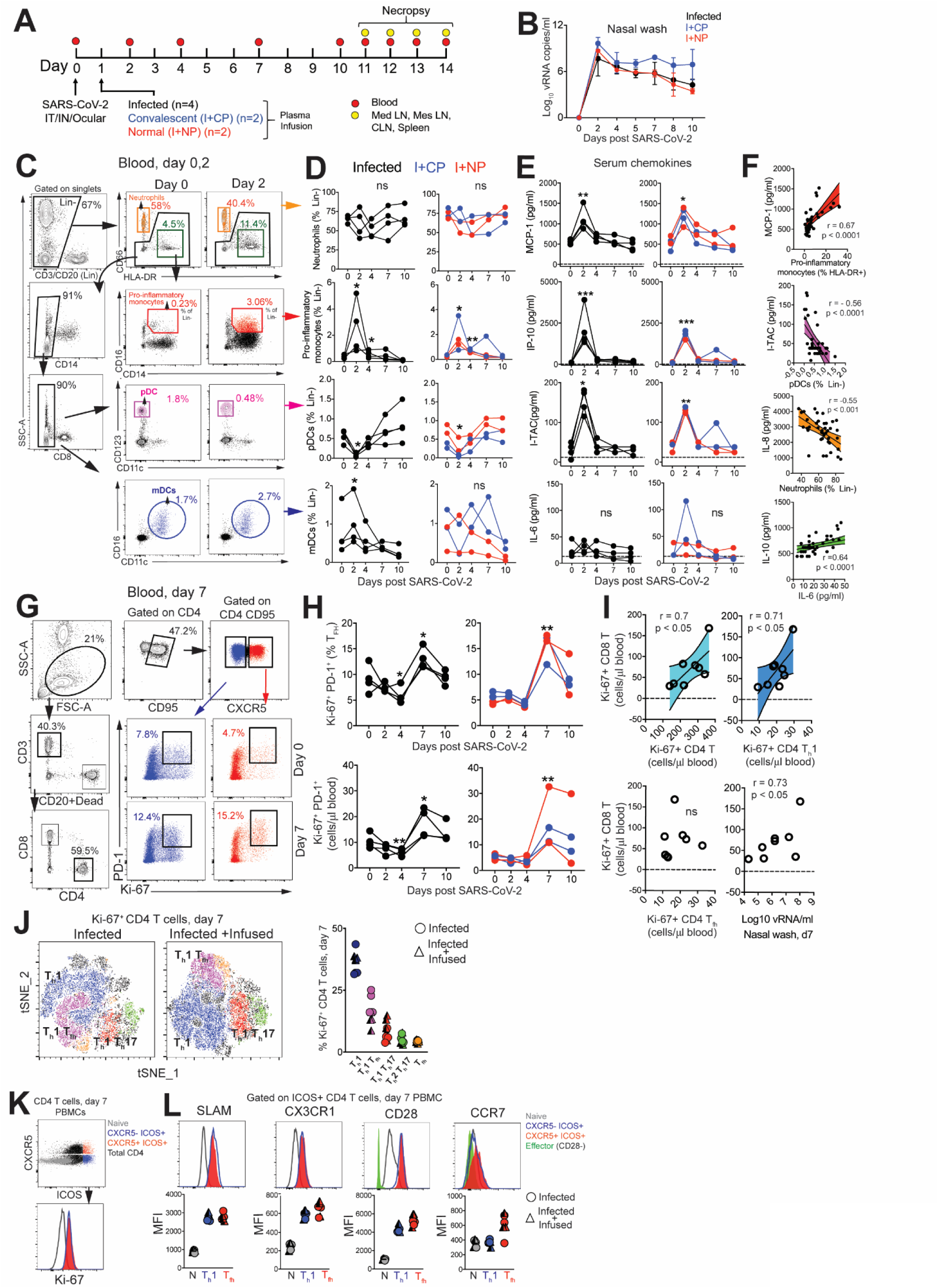
SARS-CoV-2 infection leads to a rapid and transient shift in innate immune responses and increases the number CD4 T follicular helper cells in peripheral blood. **(A)** Experimental design. Indian-origin rhesus macaques were inoculated with SARS-CoV-2 (SARS-CoV-2/human/USA/CA-CZB-59×002/2020) via the intranasal (IN), intratracheal (IT) and ocular route. Twenty-four hours later, animals were infused with either COVID-19 convalescent human plasma (I+CP; blue symbols), or normal plasma (I+NP; red symbols) (both at 4ml/kg), and four animals did not receive any plasma (infected; black symbols). Blood was sampled over the course of infection and tissues were collected at necropsy (11-14 DPI) for immune profiling. **(B)** Mean viral RNA (+range) in each of the groups within nasal washes **(C)** Flow plot illustrating gating strategy to identify innate immune subsets in whole blood. **(D**) Kinetics of innate immune responses (*p< 0.05, **p< 0.01 relative to Day 0); Lin-corresponds to CD3-CD20-. (**E**) Kinetics of serum chemokines MCP-1, IP-10, and I-TAC (*p< 0.05, **p< 0.01, ***p< 0.001 relative to Day 0). **(F)** Correlation plot of innate immune cells against corresponding chemokines, and IL-10 vs IL-6. **(G)** Gating strategy to capture Ki-67^+^ PD-1^+^ CXCR5- and CXCR5+ CD4 T cells in whole blood. **(H)** Kinetics show frequency and absolute counts of Ki-67^+^ PD-1^+^ CD4 T_fh_ cells (*p< 0.05, **p< 0.01 relative to Day 0) (**I**) correlation plots of Ki-67+CD8 T cells against Ki-67+CD4 T cells, Ki-67+ CD4 T_h_1 cells, Ki-67+ CD4 T_fh_ cells, and nasal wash vRNA (all day 7) (**J**) tSNE plot of 16,197 CD4 Ki-67+ events at Day 7 from infected and 22,406 events from infected + infused animals; dot plot illustrates proliferating (Ki-67+) CD4 T cell subsets. (**J**) Flow plot indicating four different populations (Naive CD4 T cells, CXCR5-ICOS+, CXCR5+ ICOS+, Total CD4+) along with corresponding Ki-67 expression, shows ICOS+ CD4 T cells in cell cycle at Day 7. (**K**) Histograms and median fluorescence intensity (MFI) dot plots illustrate relative expression of SLAM, CX3CR1, CD28, and CCR7 within four different populations at Day 7.

### SARS-CoV-2 infection leads to a rapid and transient shift in innate immune responses and increases the number CD4 T follicular helper cells in peripheral blood

We first sought to understand innate immune dynamics following infection. Evaluation of innate immune cell subsets in the peripheral blood (**Figure 1C**) revealed no significant changes in either the proportion or absolute counts of neutrophils over time (**Figure 1D**). However, rapid and divergent changes in specific myeloid cell subsets were observed. While CD14+ CD16+ pro-inflammatory monocytes significantly increased at Day 2 with a corresponding increase in CX3CR1 expression (**Figure S1C**), pDCs decreased in peripheral blood. We also noted a significant increase in myeloid DCs (mDC) within the infected group. Evaluation of soluble factors showed that pro-inflammatory chemokines monocyte chemoattractant protein (MCP-1), interferon γ-induced protein-10 (IP-10), interferon inducible T cell α chemoattractant (I-TAC) were significantly elevated at Day 2 and returned to baseline levels soon thereafter (**Figure 1E**). We did not observe significant elevations in pro-inflammatory cytokines interleukin (IL)-6 (**Figure 1E**), IL-1ß, or tumor necrosis factor (TNF)α (data not shown). Analysis of peripheral blood samples revealed a direct relationship between serum MCP-1 levels and pro-inflammatory monocytes over the course of infection, while pDCs and neutrophil frequencies were inversely related to levels of I-TAC and IL-8 respectively (**Figure 1F**). While no statistically significant changes occurred with IL-6 or IL-10, the levels of both cytokines were correlated over the course of infection. Overall, the innate immune cellular dynamic in the peripheral blood of healthy adult rhesus macaques following SARS-CoV-2 infection was characterized by a rapid and transient shift with resolution of systemic inflammation early during the course of infection.

To assess the increase in CD4 T_fh_ cells attributable to SARS-CoV-2 and ascertain if T_fh_ dynamics were altered following passive immunization with CP, we profiled peripheral blood samples to capture effector T cell responses. No evidence of general lymphopenia was observed over the course of infection (**Figure S1D**). Frequency and absolute counts of activated CXCR5^+^ CD4 T_fh_ cells, identified by co-expression of Ki-67 and PD-1, significantly increased in all animals at Day 7 regardless of plasma intervention (**Fig. 1G-H**). At the apex of the effector response, Ki-67^+^ CD4 T cells, specifically the T_h_1 but not the T_fh_ subset was strongly associated with proliferating CD8 T cells (**Fig. 1I**). In turn, we observed strong antigen-dependent induction of CD8 T cells evidenced by the association between SARS-CoV-2 vRNA from nasal washes and proliferating (Ki67+) CD8 T cells.

Evaluation of infection-induced changes in CD4 T cell differentiation at Day 7 revealed a strong phenotypic shift to T_h_1 effectors (CXCR3+), T_h_1 polarized T_fh_ cells (CXCR3+ CXCR5+) and T_h_1 Th17 (CXCR3+ CCR6+) CD4 T cells (**Figure 1J, Figure S2A**). Correspondingly, the data showed accumulation of CD4 T_h_1 cells in the periphery at Day 7 (**Figure S2B**). While T_h_2 CD4 cells did not peak at Day 7, there was an increase of T_h_17 CD4 cells likely due to the mucosal nature of the infection (**Figure S2C-D**). Using the acute activation marker, inducible costimulator (ICOS), to identify proliferating (Ki67+) CD4 T cells at Day 7 (**Figure 1K)**, we found that ICOS+CXCR5- and CXCR5+ CD4 T cells subsets expressed the T_h_1 marker signaling lymphocyte adhesion molecule (SLAM) and the effector molecule CX3CR1, a marker potentially for newly generated memory CD4 T cell subsets, consistent with their activation status (**Figure 1L).** Neither the ICOS+CXCR5-nor the CXCR5+ CD4 T cell subsets downregulated CD28 and both subsets expressed CCR7 at levels comparable to or greater than naive cells, indicative of a lymph node-homing phenotype. To assess CD4 T cell functionality, cytokine production was evaluated *ex vivo* following stimulation with PMA and ionomycin. Two distinct CD4 T cells were identified – a degranulating CD107a+b subset with the majority of degranulating CD4 T cells expressing interferon gamma (IFN-γ) and TNFα but not IL-2 or IL-17; and an IL-21-producing subset (**Figure S2E**). In contrast, the majority of IL-21-expressing cells produced IL-2, IL-17 and co-produced TNFα and IFN-γ. Thus, CD4 T cell polyfunctionality was preserved during SARS-CoV-2 infection. Despite the increase in activated T_fh_ cells, levels of CXCL13 did not increase significantly following SARS-CoV-2 infection (**Figure S2F**).

### CD4 T_fh_ cells targeting the spike (S) and nucleocapsid (N) proteins are generated following SARS-CoV-2 infection

Based on the significant increase in systemic CD4 T_fh_ cells following SARS-CoV-2 exposure, we sought to understand splenic involvement during the germinal center phase of the immune response. To this end, we quantified GC T_fh_ cells in the spleen at necropsy (between Day 11 and Day 14 post infection) and compared the values to those seen in animals who had not been exposed to SARS-CoV-2. The results suggested the initiation of a GC response within the spleen following infection (**Figure S3A**). We observed the majority of the GC T_fh_ cells did not express Foxp3 indicating that GC T_fh_ cells predominated over the GC T follicular regulatory cell (T_fr_) subset (**Figure S3B**). To conclusively assess SARS-CoV-2-induced responses, we stimulated cryopreserved splenocytes with mega pools - overlapping peptides covering multiple T cell epitopes in S, N, and membrane (M) proteins, and spanning the open reading frames (ORF1,3,8) of SARS-CoV-2. PMA/Ionomycin was used as a positive control while DMSO-treated cells served as negative controls. Using activation induced marker (AIM) assay, SARS-CoV-2-specific CD4 T cells were identified based on co-expression of OX40 and CD25 (**Figure 2A**). Based on cell recovery, 2 animals were excluded from the analysis of Ag-specific responses and 3 animals were excluded from analysis of N-specific responses (**Table S2**)

**Figure 2.**
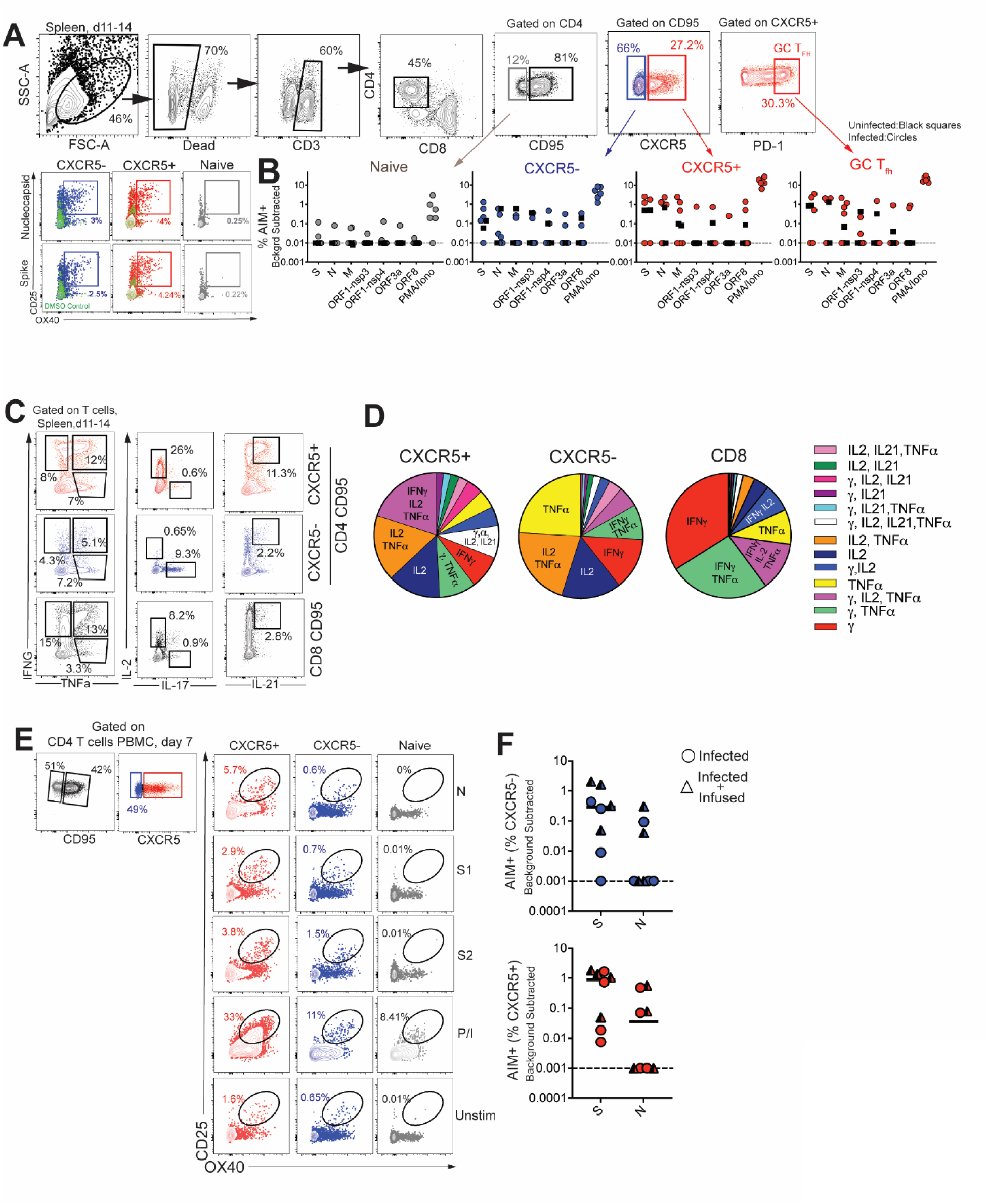
CD4 T_fh_ cells targeting the spike (S) and nucleocapsid (N) are generated following SARS-CoV-2 infection. **(A)** Gating strategy for identifying SARS-CoV-2 specific CD4 T cells in spleen following stimulation with peptide megapools **(B)** Scatter plot showing AIM+ CD4 subsets; naive, CXCR5-, CXCR5+, and CXCR5+ PD-1^++^ GC T_fh_ cells. The dashed line represents undetectable responses assigned a value of 0.01% (**C**) Cytokine profiles (IFN-γ, IL-2, TNFα, IL-17, IL-21) of CXCR5+, CXCR5-, and CD8+CD95+ T cells in spleen following PMA/Ionomycin stimulation. (**D**) Pie chart demonstrates polyfunctionality of T cell subsets following SARS-CoV-2 infection. **(E**) Gating strategy for identifying SARS-CoV-2 specific CD4 T cells in PBMCs. **(F)** AIM+ CXCR5- and CXCR5+ CD4 subsets in PBMCs at Day 7. Black squares denote SARS-CoV-2 unexposed animals. Circles denote infected and triangles denote infected+infused animals.

Following subtraction of AIM+ responses in DMSO-treated cells, CD4 T cell responses to S and N were detected in 30% of animals. Furthermore, PD-1++ GC T_fh_ cells, reactive to S, N, and M were observed indicative of SARS-CoV-2-induced GC response in the spleen (**Figure 2B**). It should be noted, however, that responses to S, N, M were also detected in unexposed animals suggestive of cross-reactive T cells to endemic coronaviruses, as has been reported in humans (13). Evaluation of CD4 T cell polyfunctionality in the spleen by ICS in response to PMA/Ionomycin stimulation revealed that CXCR5+ CD4 T cells were clearly distinguishable from CXCR5-subsets in their ability to co-produce IFN-γ, IL-2, TNFα, and IL-21. In contrast, the CXCR5-subset did not produce IL-21 yet was able to co-produce IL-2 and TNFα, or, alternatively, either IFN-γ, IL-2, or TNFα. In contrast, CD8 T cells were predominantly IFN-γ producers (**Figure 2C-D**). Together, the data from the spleen demonstrate presence of robust IL-21-producing T_fh_ population and the generation of S- and N-specific GC T_fh_ cells. Consistent with data from the spleen, antigen-specific responses against S and N were also observed in peripheral blood at Day 7 (**Figure 2E-F**). Together, these data demonstrate that S- and N-specific CD4 T_fh_ cells are elicited following SARS-CoV-2 infection.

### SARS-CoV-2 infection induces germinal center responses in mediastinal lymph nodes

Having established that SARS-CoV-2 stimulates the production of CD4 T_fh_ cells, we next sought to understand whether effector T_h_1 CD4 T cells were induced in the lung. Subsequent to collagenase digestion, single cell suspensions isolated from the lung were stained with a panel of markers to delineate activated CD4 T cells. Based on cell recovery and events following acquisition, data from 5 out of 8 animals were analyzed. We evaluated expression of Granzyme B and PD-1, both antigen-induced activation markers; mucosal homing receptors α_4_ß_7_, CCR6, and the Th1 receptor CXCR3 within CD69+ and CD69-CD4 T cell subsets (**Figure 3A**). Expression pattern of Granzyme B, PD-1, and CXCR3 in lung CD4 T cells was indicative of a T_h_1 effector CD4 response (**Figure 3B**). Furthermore, histopathology of the lung showed development of Bronchus-associated lymphoid tissue (**Figure S4A**), providing a strong rationale to assess the draining mediastinal lymph node for germinal center responses. Gross examination of the mediastinal lymph nodes was consistent with lymphadenopathy (**Figure S4B**). Based on reports detecting SARS-CoV-2 vRNA in the intestine (24), the mesenteric lymph nodes were assessed as a secondary site of viral dissemination. Following isolation of single-cell suspensions after collagenase digestion, cells from the mediastinal lymph nodes, cervical lymph nodes, and mesenteric lymph nodes were stained with a panel of markers to define GC T_fh_ cells, GC B cells, and follicular dendritic cells (FDCs). As illustrated, mediastinal lymph nodes showed a distinct CXCR5+ PD-1++ GC T_fh_ subset and Bcl-6+ CD20+ GC B cells (**Figure 3C)**. FDCs were identified based on expression of the complement receptor CD21 (clone B-Ly4; **Figure S4C**), instrumental in immune complex trapping, within the CD45- CD3- CD20- cell population (28). The number of FDCs strongly correlated with the frequencies of both GC B cells and GC T_fh_ cells (**Figure S4D**).

**Figure 3.**
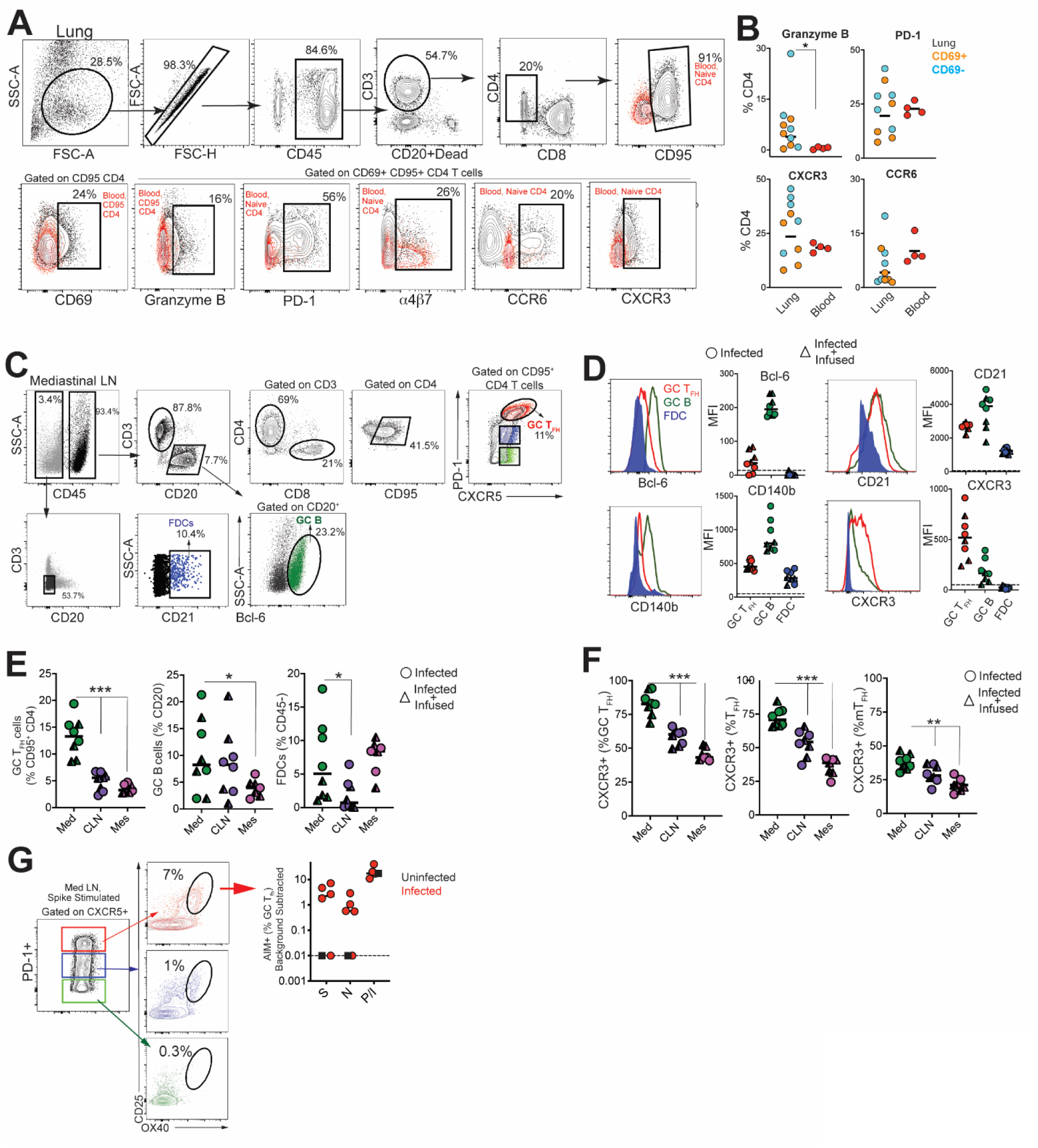
SARS-CoV-2 infection induces germinal center responses in mediastinal lymph nodes. **(A)** Gating strategy for identifying CD4 T cells in lung; red overlay represents paired CD4 subset from blood (either CD95- (naive) or CD95+ as indicated). **(B)** Scatter plot shows expression of Granzyme B, PD-1, CXCR3, CCR6 on CD69- and CD69+ subsets in lung and CD95+ CD4 T cells in blood. **(C)** Gating strategy for identification of GC T_fh_ cells, GC B cells and FDCs. **(D)** Relative expression of Bcl-6, CD21, CD140b, and CXCR3 within GC cell subsets. **(E)** Frequency of GC T_fh_ cells, GC B cells, FDCs significantly higher in mediastinal lymph node (*p< 0.05, ***p< 0.001). **(F)** Relative expression of CXCR3 shows majority of GC T_fh_ cells in mediastinal lymph nodes express CXCR3 (**p< 001; ***p< 0.001). (**G**) Flow plot of PD-1+ CXCR5+ (GC) T_fh_ cells show positivity for markers CD25 and OX40 following stimulation with spike; scatter plot shows specificity of GC T_fh_ cells to various SARS-CoV-2 proteins (Spike, Nucleocapsid, Membrane). The dashed line represents undetectable responses assigned a value of 0.01%. Black squares denote SARS-CoV-2 unexposed animals. Circles denote infected animals and triangles denote infected+ infused animals.

Quantifying the expression of canonical GC markers showed that Bcl-6 was exclusively expressed by GC B cells and to a lesser extent by GC T_fh_ cells (**Figure 3D**). FDC markers CD21 and platelet-derived growth factor receptor b (CD140b) (29) were also expressed by GC Tfh and B cells (**Figure 3D**). An increase in expression of the T_h_1-chemokine receptor, CXCR3, on GC T_fh_ cells was consistent with the phenotype of cells responding to viral infection. While GC B cells displayed heterogeneity in CXCR3 expression, FDCs were uniformly negative for this marker. The increased number of GC T_fh_, GC B cells and FDCs (**Figure E**) as well as the higher relative expression of CXCR3 in mediastinal lymph nodes compared to cervical and mesenteric lymph nodes indicated an active immune response to viral infection (**Figure 3F**). Consistently, we observed SARS-specific responses by GC T_fh_ cells in the mediastinal lymph node (**Figure 3G, Figure S4E).**

### Humoral responses to SARS-CoV-2 are dominated by IgG antibodies

Studies in humans have shown that greater than 90% of SARS-CoV-2 patients develop binding antibodies to S antigen within 10 days of symptom onset (30, 31). However, the kinetics of the early antibody response to S and N proteins and the contributing antibody isotypes, specifically in the setting of mild or asymptomatic clinical illness, are not well-defined. Here, we quantified concentrations of serum antibodies to S1, S2, and N antigens, and used a secondary antibody specific for macaque IgG to distinguish *de novo* IgG antibodies from passively infused human CP IgG antibodies. The data showed IgM and IgG seroconversion to S1 and S2 proteins in all animals by day 7 post-infection, with the exception of one CP animal (**Figure 4A & B**). This is consistent with reports that S- or RBD-specific IgG and IgM antibodies often appear simultaneously in blood of most humans infected with 2002 SARS or CoV-2 (30-32). Antibody responses to the N protein in humans are reported to increase 10 days following disease onset (33, 34), and interestingly, N-specific IgG was evident in all macaques by day 7 but N-specific IgM was not increased significantly (3-fold over baseline values) until day 10 in most animals. In addition, 50% of the animals failed to demonstrate a significant IgA response to all SARS CoV-2 proteins within 10 days of infection **(Figure 4C**). However, we should note that analysis of some necropsy sera suggested that IgA antibodies continue to increase after day 10 (data not shown).

**Figure 4.**
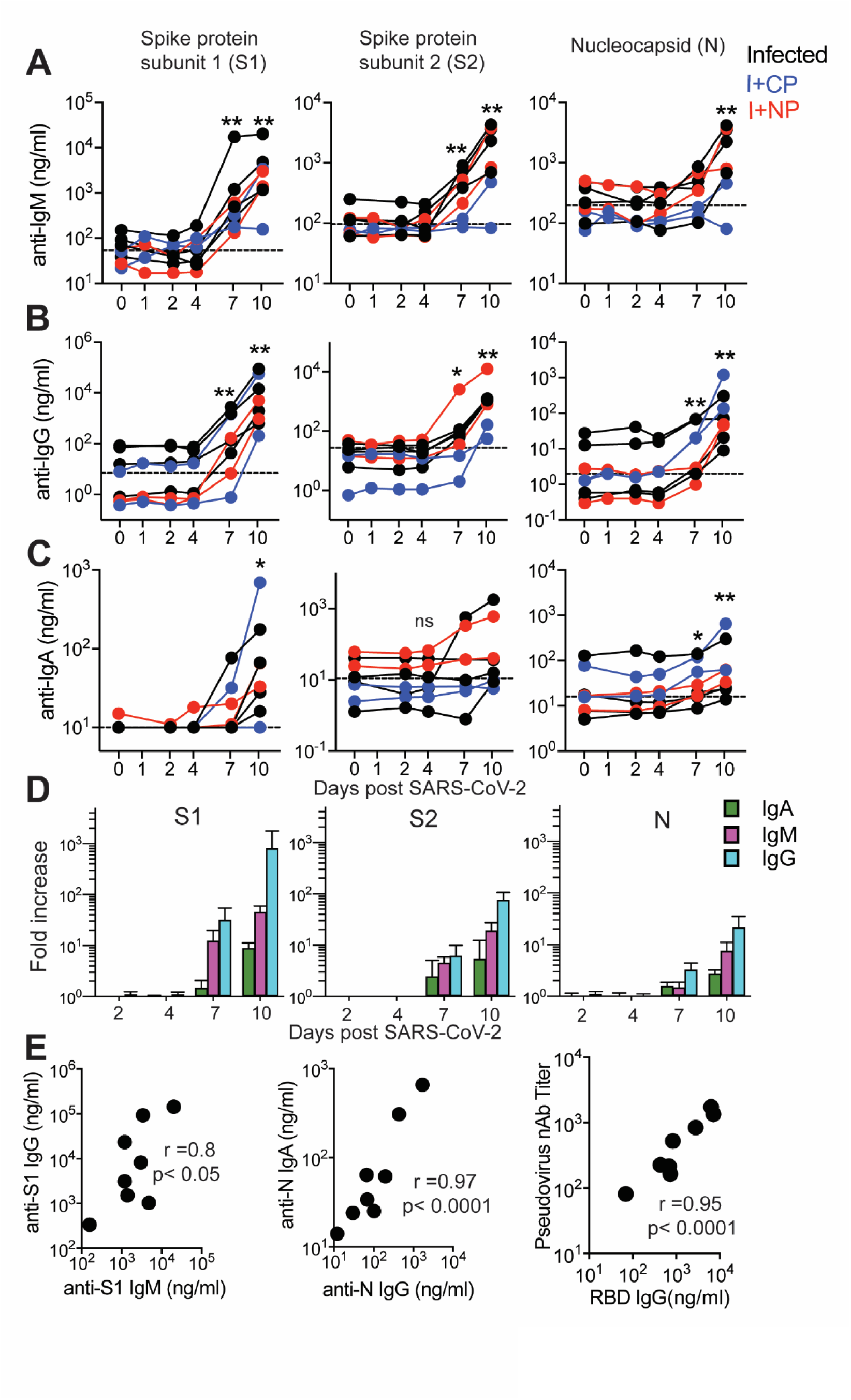
Humoral responses to SARS-CoV-2 are dominated by IgG antibodies. Concentrations of **(A)** IgM, **(B)** IgG, and **(C)** IgA antibodies specific for S1, S2, and N proteins were measured by BAMA or ELISA in serum of macaques infused with human COVID-19 convalescent plasma (CP; blue symbols) or naive plasma (NP; red symbols) and control non-infused animals (black symbols). The dashed line represents the median pre-infection (day 0) concentration for all animals. **(D)** The magnitude of the IgM, IgG and IgA antibody responses in animals that were not given human convalescent plasma was determined by dividing post-infection concentrations by those measured on day 0 in each animal. Geometric mean fold increases with SEM are shown. **(E)** Correlations between day 10 levels of S1-specific IgG and IgM, N-specific IgA and IgG, and pseudovirus neutralizing antibody titers and anti-RBD IgG antibodies measured by ELISA.

Evaluation of the magnitude of post-infection antibody responses in animals that did not receive CP plasma clearly indicated that IgG dominated the humoral response to all SARS CoV-2 proteins **(Figure 4D)**. On day 10, we observed strong correlations between S1-specific IgG and IgM and between N-specific IgA and IgG (**Figure 4E**). Also consistent with reports in infected humans, we observed a strong correlation between neutralization antibody titers and concentrations of anti-RBD IgG antibodies on day 10 (**Figure 4E**). Together, these data show rapid development of binding and neutralizing antibodies following SARS-CoV-2 infection in the context of mild or absent clinical symptoms. The appearance of antiviral IgG antibodies by day 7 with delayed induction of IgA responses suggests that early class-switching occurs after SARS-CoV-2 infection and is likely promoted by T_h_1-type T_fh_ cells.

## DISCUSSION

The importance of CD4 T_fh_ cells in the generation of plasma cells, critical for persistent antibody, places a premium on understanding the T_fh_ and GC response following SARS-CoV-2 infection. The present study adds to our understanding of immune responses to SARS-CoV-2 in three significant ways. First, we demonstrate that robust T_h_1-T_fh_ responses are observed following SARS-CoV-2 infection. Second, T_fh_ responses focused on S and N are seen within lymph nodes, circulate through peripheral blood, potentially seeding the spleen. Lastly, we show that acute antibody kinetics are characterized by induction of IgG, predominantly to S1, indicative of early class switching. Taken together, these data demonstrate that productive T_fh_ responses are elicited following SARS-CoV-2 infection in healthy adult rhesus macaques.

The innate immune dynamics induced following SARS-CoV-2 infection was characterized by a distinct increase in CD14+ CD16+ pro-inflammatory monocytes with corresponding upregulation of CX3CR1 at Day 2 likely reflective of their increased trafficking to the lower airways in response to CCR2/CCR5 ligands (35) (36-38). An increase in pro-inflammatory monocytes is observed in COVID-19 patients positively correlating with disease severity. In contrast to monocytes, the rapid depletion of pDCs in peripheral blood is consistent with their recruitment to the lower-airways where pDC-mediated type I interferon is a critical facet of the anti-viral response (39). The brisk and transient innate immune dynamics following exposure to SARS-CoV-2 are consistent with the minimal changes observed in body weight and oxygen saturation levels and mild overall disease pathology in animals.

Several studies have examined the kinetics of antibody responses in humans after the onset of symptoms and three unifying themes emerge from these data. First, in the majority of patients, antibodies to RBD of the S1 subunit are induced between 8-10 days of symptom onset, and levels of these antibodies correlate strongly with neutralizing titers (30, 31, 40). Second, plasma from the majority of COVID-19 convalescent patients does not contain high levels of neutralizing activity (41). Third, plasma antibodies in infected individuals that do develop neutralizing antibodies are minimally mutated (42). These data suggest that CD8 T cells may contribute to the control of SARS-CoV-2, and while a protracted germinal center response may not be critical for the generation of neutralizing antibodies it could improve antibody durability by enhancing plasma cell numbers. Our data add to this developing narrative by showing that in the setting of mild/asymptomatic illness, antibody responses are generated and characterized by the predominance of IgG. Intriguingly, we observed that IgM and IgG antibodies to the S1, S2 and N proteins were produced concurrently.

While it remains unknown whether immune responses elicited when naturally infected by SARS-CoV-2 will protect from re-infection, studies in rhesus macaques show that infection does protect against re-challenge, 28-35 days post first infection, signifying that some degree of a protective immune response follows infection (27). In this context, the finding that infection induces CD4 helper responses targeting major structural proteins on the virus suggests that infection is capable of producing effective CD4 help for CD8 T cells and antibody responses. Indeed, antibodies against the RBD region in S1 are elicited in the vast majority of COVID-19 patients along with robust CD4 T cell responses(31). Our data show that spike epitopes are immunogenic to both T and B cells and suggests that induction of these responses by vaccines may confer protection. While our discovery of N- and S-specific CD4 T cells in the spleen is intriguing at present we are unable to distinguish whether these cells represent cells that are seeded from circulation or are elicited *de novo* via trafficked antigen. Further studies are needed to tease apart the possibilities as this is central to understanding determinants of protective immunity. In sum, the data suggest that vaccine platforms inducing T_h_1 CD4 helper and T_fh_ helper responses are likely to succeed in eliciting robust CD8 T cell and antibody responses against SARS-CoV-2.

Recent data suggest benefit of CP therapy, in conjunction with antiviral and immunomodulators, in treating moderate to severe COVID-19 (43-45), but that CP infusion early during the course of infection may be more beneficial as antibody responses are generated within 2 weeks of symptom onset (46). Our studies, however, demonstrate that CP infusion did not abate a nascent infection. While the pooled CP had neutralization titers of 1:1149, we failed to achieve detectable neutralizing antibodies after infusion explaining the lack of efficacy. These results highlight the prerequisite for convalescent plasma to have sufficiently high titers of neutralizing antibodies to treat COVID-19 patients. More controlled studies are required to confidently determine whether CP achieves neutralizing titers that can protect either prophylactically or therapeutically. Data supporting this would have considerable potential not only for therapeutics but also the efficacy of vaccines.

In summary, the current study adds to our understanding of the CD4 helper responses to SARS-CoV-2 infection and provides an important foundation for harnessing the mechanisms that stimulate robust CD4 T_fh_ responses in the context of an effective vaccine.

## Supporting information

Animal Information

Spleen AIM Assay conditions

Mediastinal LN AIM Assay conditions

Clinical symptoms and complete blood counts following SARS-CoV-2 infection

SARS-CoV-2 infection elicits Ki67+PD-1+ Th1 cells at Day 7 that are polyfunctional

Splenic GC TFH, Tfr, and Treg populations during SARS-CoV-2 infection

SARS-CoV-2 infection increases mediastinal lymph node size and peptide pool stimulation affect on CXCR5+, CXCR5- and Naive T cells.

## ACKNOWLEDGEMENTS

The authors are grateful to Lourdes Adamson and Nicole Drazenovich for BSL3 training and facilitating access to CIID BSL3. The authors are grateful to Greg Hodges for facilitating the animal experiments in CNPRC ABSL3. We are extremely grateful to the primate center staff Wilhelm Von Morgenland, Miles Christensen, David Bennet, Vanessa Bakula, James Schulte, Jose Montoya, Joshua Holbrook, John McAnelly, Christopher Nelson, and David Bennett for animal sampling in ABSL3 at the CNPRC. Mark Allen provided necropsy technical expertise in the BSL-3 necropsy suite. The authors are grateful to Amanda Carpenter, Peter Nham, and Bryson Halley at the Primate Assay Laboratory Core. We thank Jeffrey Roberts and the CNPRC leadership for their support in facilitating these experiments and acknowledge assay resources provided by the Primate Assay Laboratory Core and CNPRC base grant. The 10F12 anti-monkey IgA monoclonal antibody was obtained from the NIH Nonhuman Primate Reagent Resource supported by AI126683 and OD010976. The study was funded by internal seed grants to CNRPC and CIID, R21 AI143454-02S1 (SSI), FAST GRANT-George Mason University (SSI), Animal Models of Infectious Diseases Training Program T32AI060555 (SRE) and National Institutes of Health contract Nr. 75N9301900065 (A.S. and D.W).

## METHODS

### Rhesus Macaques

Eight colony-bred Indian origin rhesus macaques (*Macaca mulatta*) were housed at the California National Primate Research Center and maintained in accordance with American Association for Accreditation of Laboratory Animal Care guidelines and Animal Welfare Act/Guide. As described by us (47), strict social distancing measures were implemented at the CNPRC at the start of the pandemic in March to reduce risk of human-to-rhesus SARS-CoV-2 transmission. Animals were screened for SARS-CoV-2 and housed in barrier rooms with increased PPE requirements prior to study assignment. Animals were four to five years of age with a median weight of 8.6 kg (range 5.4-10.7 kg), were SIV- STLV- SRV-. Animals were seronegative for SARS-CoV-2 at study initiation. Sex distribution within experimental groups was as follows; Infected (n=3 females, n=1 male); Infected + Convalescent Plasma (n=2 males); Infected + Normal Plasma (n=2 males). Table S1 provides details of the animals in the study. For blood collection, animals were anesthetized with 10 mg/kg ketamine hydrochloride injected i.m. For virus inoculation and nasal secretion sample collection, animals were additionally anesthetized with 15-30 ug/kg dexmedetomidine HCl inject i.m. and anesthesia was reversed with 0.07-0.15 mg/kg atipamezole HCl injected i.m.

### Virus and inoculations

SARS-CoV-2 virus (2019-nCoV/USA-CA9/2020) was isolated from the nasal swab of a COVID-19 patient with acute respiratory distress syndrome admitted to University of California, Davis Medical Center, Sacramento (48). Vero cells (ATCC CCL-81) were used for viral isolation and stock expansion. The passage 2 viral stock (SARS-CoV-2/human/USA/CA-CZB-59×002/2020) used for animal inoculations had a titer of 1.2×10^6^ PFU/ml (Genbank accession number: MT394528). To recapitulate relevant transmission routes of SARS-CoV-2, animals were inoculated with 1 ml stock instilled into the trachea, 1 ml dripped intranasally, and a drop of virus stock in each conjunctiva.

### Convalescent plasma and infusions

Convalescent plasma was sourced from Vitalant and represented a pool from up to four donors. Plasma was pooled prior to infusion into monkeys. Pooled plasma had a nAb titer of 1:1149, binding antibody titers for SARS-CoV-2 antigen were as follows; anti-S1-IgG, 24.5 µg/ml; anti-S2 IgG, 2.9 µg/ml; anti-NC IgG, 10.7 µg/ml. Normal plasma was collected prior to the COVID-19 pandemic and was negative for SARS-Cov2 antibody. Concentrations were as follows; anti-S1-IgG, 0.00004µg/ml; anti-S2 IgG, 0.003 µg/ml; anti-N IgG, 0.001 µg/ml. Twenty fours following virus inoculation, animals were infused with plasma at 4 ml/kg volume (total volume infused was 33-39 ml) at an infusion rate of 1 ml/minute. Control animals (n=4) were not infused.

### Specimen collection and processing

On days 2, 4, 5, 7, 8, and 10, a 5-french feeding tube was inserted into the nasal cavity. 2 ml PBS were instilled through each nostril and the maximum volume was aspirated, secretions were spun at 800 g for 10 min and 1 ml of supernatant and cell pellet were lysed in 3 ml Trizol LS for RNA isolation. EDTA-anticoagulated blood was collected on Days 0, 2, 4, 7, and 10 for immunophenotyping. PBMCs were isolated from whole blood collected in CPT vacutainer tubes, sampled at Day 7 and necropsy, by density gradient centrifugation as previously described (49). For serum, coagulated blood was centrifuged at 800 g for 10 min to pellet clotted cells, followed by extraction of supernatant and storage at −80°C. Lymph nodes, spleen, and lung tissue were obtained at necropsy and digested enzymatically using collagenase followed by manual disruption to obtain single cell suspensions for flow cytometry based assays.

### Activation induced Marker (AIM) assay

Cells were stimulated with overlapping peptide pools representing SARS-CoV-2 and responding cells were identified by upregulation of activation markers, as described previously (49, 50). All antigens were used at a final concentration of 2 μg/mL in a stimulation cocktail made with using 0.2 μg of CD28 and 0.2 μg CD49d costimulatory antibodies per test. Unstimulated controls were treated with volume-controlled DMSO (Sigma-Aldrich). Tubes were incubated in 5% CO_2_ at 37°C overnight. Following an 18 h stimulation, the cells were stained, fixed, and acquired the same day. AIM assays on splenocytes and mediastinal lymph nodes were performed on cryopreserved cells (Table S2). AIM assay on day 7 PBMCs were performed on fresh cells. Phenotype panel on LNs and PBMCs was performed using standard flow cytometry assays.

### vRNA quantitation by quantitative real time polymerase chain reaction (qRT-PCR)

Trizol lysed nasal samples were processed using a modified Qiagen RNeasy Lipid Tissue Mini Kit protocol. Briefly, 5.6ul polyacryl carrier was added to trizol lysate, followed by 1/10 volume BCP and phase separated as described in Qiagen protocol. 8ul of eluted RNA was DNase treated with ezDNase per kit instructions and converted to cDNA with Superscript IV using random primers in a 20ul reaction and quantified in quadruplicate by qPCR on an Applied Biosystems QuantStudio 12K Flex Real-Time PCR System using Qiagen QuantiTect Probe PCR Mastermix with primers and probe that target the SARS-CoV-2 Nucleocapsid (forward 5’-GTTTGGTGGACCCTCAGATT-3’, reverse 5’-GGTGAACCAAGACGCAGTAT-3’, probe 5’-/5-FAM/TAACCAGAA/ZEN/TGGAGAACGCAGTGGG/3IABkFQ/-3’).

### Serum cytokines

Serum cytokines. Luminex® (NHP Cytokine Luminex Performance Pre-mixed kit, R&D, FCSTM21) was performed to evaluate cytokines in rhesus macaque sera. The assay was performed according to the manufacturer’s protocol. The beads for each sample, control, and standard curve point were interrogated in a Luminex® 200 dual laser instrument (Luminex, Austin, TX) which monitors the spectral properties of the beads and amount of associated phycoerythrin (PE) fluorescence for each sample. xPONENT® software was used to calculate the median fluorescent index and calculate the concentration for each cytokine in

### Flow cytometry

Cell staining was performed as previously described. Whole blood and single cell suspensions from the lung and lymph nodes were stained fresh and acquired the same day. Staining on spleen was performed on cryopreserved samples. Fluorescence was measured using a BD Biosciences FACSymphony™ with FACSDiva™ version 8.0.1 software. Compensation, gating and analysis were performed using FlowJo (Version 10). Polyfunctionality plots were generated using SPICE (v 6) (51). Reagents used for flow cytometry are listed in Table S2.

### BAMA for IgG and IgM antibodies to S1, S2 and N proteins

A customized BAMA was developed to simultaneously measure antibodies to the following recombinant SARS CoV-2 proteins (all from SinoBiologicals, Wayne, PA): S1 (#40591-V08H), S2 extracellular domain (#40590-V08B) and nucleocapsid (N; #40588-V08B). Briefly, proteins were dialyzed in PBS and conjugated to Bioplex Pro carboxylated magnetic beads (BioRad, Hercules, CA) as previously described (49). Standard and serum samples treated with 1% TritonX-100 detergent were serially diluted in PBS containing 1% BSA, 0.05% azide, and 0.05% Tween-20 and mixed with beads overnight at 1100rpm and 4°C. The following day, the beads were washed and treated with biotinylated antibody followed by neutralite avidin-phycoerythrin (Southern Biotechnology Associates: SBA, Birmingham, AL) as described (Phillips 2017). A BioRad Bioplex 200 and BioManager software were used to measure fluorescent intensity and construct standard curves for interpolation of antibody concentrations in test samples.

The standard was pooled serum from macaques infected for 11-14 days with SARS CoV-2. The following humanized (IgG1) monoclonal antibodies were used to estimate concentrations of IgG, IgM, and IgA antibodies in the rhesus serum standard: anti-S1 RBD (Genscript #HC2001), anti-S2 (SinoBiologicals #40590-D001) and anti-NC (Genscript #HC2003). Human and rhesus IgG antibodies were both detected in these calibration assays using biotinylated affinity-purified goat anti-human IgG γ chain polyclonal antibody (SBA #2048-08). In subsequent BAMA assays for SARS CoV-2-specific rhesus macaque IgG antibodies, biotinylated mouse anti-monkey IgG γ chain monoclonal antibody (SBA cat#4700-08) was used as the secondary antibody. IgM antibodies were detected using biotinylated affinity-purified goat anti-human IgM µ chain polyclonal antibody (SBA#2020-08) which cross-reacts well with macaque IgM. Results obtained for IgM in the rhesus standard were multiplied by 3.3 to account for under-estimation by the monomeric IgG monoclonal antibody standard. Macaque IgA antibodies were detected with the antibodies described below. IgG, IgM or IgA antibodies had to be increased 3-fold over the day 0 value to be considered significant

### ELISA for SARS-specific IgA and antibodies to RBD

These assays were done using methods similar to those described (52) and Immulon 4 microtiter plates (VWR, Radnor, PA) coated with 100ng per well of S1, S2, N or RBD protein (SinoBiological #40592-VNAH). For IgA assays, a pooled rhesus serum collected day 14 after infection with SARS CoV-2 was used as standard after depletion of IgG using GE Healthcare Protein G Sepharose (Sigma, St. Louis, MO) as described (52). Test sera were also depleted of IgG to facilitate detection of low levels of IgA antibodies. Macaque IgA was detected using a mixture of biotinylated clone 10F12 (NHP Reagent Resource) and biotinylated clone 40.15.5 (Ward et al 1995) anti-rhesus IgA monoclonal antibodies, which do not cross-react with human IgA and, when combined, appear to recognize all allotypes of rhesus macaque IgA (Kozlowski, personal observation). For RBD IgG assays, the above pooled rhesus serum standard and secondary monoclonal antibody specific for macaque IgG were used.

### Neutralizing assay

Pseudovirus neutralization assay was performed as described (53).

### Statistics

Statistical analyses were performed using GraphPad Prism 8.4.2. Within group comparisons, such as immune responses and antibody levels at different time points, were done using the two-tailed Wilcoxon matched-pairs signed rank test. For correlation analysis, the two-tailed Spearman rank correlation test was used.

## References

1. WHO announces COVID-19 outbreak a pandemic. World Health Organization (2020). 2020.

2. Available from: https://coronavirus.jhu.edu/map.html.

3. Huang C, Wang Y, Li X, Ren L, Zhao J, Hu Y, Zhang L, Fan G, Xu J, Gu X, Cheng Z, Yu T, Xia J, Wei Y, Wu W, Xie X, Yin W, Li H, Liu M, Xiao Y, Gao H, Guo L, Xie J, Wang G, Jiang R, Gao Z, Jin Q, Wang J, Cao B. Clinical features of patients infected with 2019 novel coronavirus in Wuhan, China. Lancet. 2020;395(10223):497-506. Epub 2020/01/28. doi: 10.1016/S0140-6736(20)30183-5. PubMed PMID: 31986264; PMCID: PMC7159299.

4. Corey L, Mascola JR, Fauci AS, Collins FS. A strategic approach to COVID-19 vaccine R&D. Science. 2020;368(6494):948-50. Epub 2020/05/13. doi: 10.1126/science.abc5312. PubMed PMID: 32393526.

5. Graham BS. Rapid COVID-19 vaccine development. Science. 2020;368(6494):945-6. Epub 2020/05/10. doi: 10.1126/science.abb8923. PubMed PMID: 32385100.

6. Amanat F, Krammer F. SARS-CoV-2 Vaccines: Status Report. Immunity. 2020;52(4):583-9. Epub 2020/04/08. doi: 10.1016/j.immuni.2020.03.007. PubMed PMID: 32259480; PMCID: PMC7136867.

7. Pulendran B, Ahmed R. Immunological mechanisms of vaccination. Nat Immunol. 2011;12(6):509-17. Epub 2011/07/09. doi: 10.1038/ni.2039. PubMed PMID: 21739679; PMCID: PMC3253344.

8. Amanna IJ, Carlson NE, Slifka MK. Duration of humoral immunity to common viral and vaccine antigens. N Engl J Med. 2007;357(19):1903-15. Epub 2007/11/09. doi: 10.1056/NEJMoa066092. PubMed PMID: 17989383.

9. Renhong Yan YZ, Yaning Li3, Lu Xia, Yingying Guo 1,2, Qiang Zhou. Structural basis for the recognition of the SARS-CoV-2 by full-length human ACE2. Science. 2020. doi: Structural basis for the recognition of the SARS-CoV-2 by full-length human ACE2.

10. Bentebibel SE, Khurana S, Schmitt N, Kurup P, Mueller C, Obermoser G, Palucka AK, Albrecht RA, Garcia-Sastre A, Golding H, Ueno H. ICOS(+)PD-1(+)CXCR3(+) T follicular helper cells contribute to the generation of high-avidity antibodies following influenza vaccination. Sci Rep. 2016;6:26494. doi: 10.1038/srep26494. PubMed PMID: 27231124; PMCID: PMC4882544.

11. Crotty S. T Follicular Helper Cell Biology: A Decade of Discovery and Diseases. Immunity. 2019;50(5):1132-48. Epub 2019/05/23. doi: 10.1016/j.immuni.2019.04.011. PubMed PMID: 31117010; PMCID: PMC6532429.

12. Iyer SS, Gangadhara S, Victor B, Gomez R, Basu R, Hong JJ, Labranche C, Montefiori DC, Villinger F, Moss B, Amara RR. Codelivery of Envelope Protein in Alum with MVA Vaccine Induces CXCR3-Biased CXCR5+ and CXCR5-CD4 T Cell Responses in Rhesus Macaques. J Immunol. 2015;195(3):994–1005. doi: 10.4049/jimmunol.1500083. PubMed PMID: 26116502; PMCID: PMC4506863.

13. Grifoni A, Weiskopf D, Ramirez SI, Mateus J, Dan JM, Moderbacher CR, Rawlings SA, Sutherland A, Premkumar L, Jadi RS, Marrama D, de Silva AM, Frazier A, Carlin AF, Greenbaum JA, Peters B, Krammer F, Smith DM, Crotty S, Sette A. Targets of T Cell Responses to SARS-CoV-2 Coronavirus in Humans with COVID-19 Disease and Unexposed Individuals. Cell. 2020. Epub 2020/05/31. doi: 10.1016/j.cell.2020.05.015. PubMed PMID: 32473127; PMCID: PMC7237901.

14. Sekine T, Perez-Potti A, Rivera-Ballesteros O, Strålin K, Gorin J-B, Olsson A, Llewellyn-Lacey S, Kamal H, Bogdanovic G, Muschiol S, Wullimann DJ, Kammann T, Emgård J, Parrot T, Folkesson E, Rooyackers O, Eriksson LI, Sönnerborg A, Allander T, Albert J, Nielsen M, Klingström J, Gredmark-Russ S, Björkström NK, Sandberg JK, Price DA, Ljunggren H-G, Aleman S, Buggert M. Robust T cell immunity in convalescent individuals with asymptomatic or mild COVID-19. bioRxiv. 2020:2020.06.29.174888. doi: 10.1101/2020.06.29.174888.

15. Wang YD, Sin WY, Xu GB, Yang HH, Wong TY, Pang XW, He XY, Zhang HG, Ng JN, Cheng CS, Yu J, Meng L, Yang RF, Lai ST, Guo ZH, Xie Y, Chen WF. T-cell epitopes in severe acute respiratory syndrome (SARS) coronavirus spike protein elicit a specific T-cell immune response in patients who recover from SARS. J Virol. 2004;78(11):5612-8. Epub 2004/05/14. doi: 10.1128/JVI.78.11.5612-5618.2004. PubMed PMID: 15140958; PMCID: PMC415819.

16. Li CK, Wu H, Yan H, Ma S, Wang L, Zhang M, Tang X, Temperton NJ, Weiss RA, Brenchley JM, Douek DC, Mongkolsapaya J, Tran BH, Lin CL, Screaton GR, Hou JL, McMichael AJ, Xu XN. T cell responses to whole SARS coronavirus in humans. J Immunol. 2008;181(8):5490-500. Epub 2008/10/04. doi: 10.4049/jimmunol.181.8.5490. PubMed PMID: 18832706; PMCID: PMC2683413.

17. Kuri-Cervantes L, Pampena MB, Meng W, Rosenfeld AM, Ittner CAG, Weisman AR, Agyekum R, Mathew D, Baxter AE, Vella L, Kuthuru O, Apostolidis S, Bershaw L, Dougherty J, Greenplate AR, Pattekar A, Kim J, Han N, Gouma S, Weirick ME, Arevalo CP, Bolton MJ, Goodwin EC, Anderson EM, Hensley SE, Jones TK, Mangalmurti NS, Luning Prak ET, Wherry EJ, Meyer NJ, Betts MR. Immunologic perturbations in severe COVID-19/SARS-CoV-2 infection. bioRxiv. 2020:2020.05.18.101717. doi: 10.1101/2020.05.18.101717.

18. Laing AG, Lorenc A, Del Molino Del Barrio I, Das A, Fish M, Monin L, Munoz-Ruiz M, Mckenzie D, Hayday T, Francos Quijorna I, Kamdar S, Joseph M, Davies D, Davis R, Jennings A, Zlatareva I, Vantourout P, Wu Y, Sofra V, Cano F, Greco M, Theodoridis E, Freedman J, Gee S, Nuo En C, Julie, Ryan S, Bugallo Blanco E, Peterson P, Kisand K, Haljasmagi L, Martinez L, Merrick B, Bisnauthsing K, Brooks K, Ibrahim M, Mason J, Lopez Gomez F, Babalola K, Abdul-Jawad S, Cason J, Mant C, Doores K, Seow J, Graham C, di Rosa F, Edgeworth J, Shankar Hari M, Hayday A. A consensus Covid-19 immune signature combines immuno-protection with discrete sepsis-like traits associated with poor prognosis. medRxiv. 2020:2020.06.08.20125112. doi: 10.1101/2020.06.08.20125112.

19. Sterlin D, Mathian A, Miyara M, Mohr A, Anna F, Claer L, Quentric P, Fadlallah J, Ghillani P, Gunn C, Hockett R, Mudumba S, Guihot A, Luyt C-E, Mayaux J, Beurton A, Fourati S, Lacorte J-M, Yssel H, Parizot C, Dorgham K, Charneau P, Amoura Z, Gorochov G. IgA dominates the early neutralizing antibody response to SARS-CoV-2. medRxiv. 2020:2020.06.10.20126532. doi: 10.1101/2020.06.10.20126532.

20. Mathew D, Giles JR, Baxter AE, Greenplate AR, Wu JE, Alanio C, Oldridge DA, Kuri-Cervantes L, Pampena MB, D’Andrea K, Manne S, Chen Z, Huang YJ, Reilly JP, Weisman AR, Ittner CAG, Kuthuru O, Dougherty J, Nzingha K, Han N, Kim J, Pattekar A, Goodwin EC, Anderson EM, Weirick ME, Gouma S, Arevalo CP, Bolton MJ, Chen F, Lacey SF, Hensley SE, Apostolidis S, Huang AC, Vella LA, Betts MR, Meyer NJ, Wherry EJ. Deep immune profiling of COVID-19 patients reveals patient heterogeneity and distinct immunotypes with implications for therapeutic interventions. bioRxiv. 2020:2020.05.20.106401. doi: 10.1101/2020.05.20.106401.

21. Neidleman J, Luo X, Frouard J, Xie G, Gurjot G, Stein ES, McGregor M, Ma T, George AF, Kosters A, Greene WC, Vasquez J, Ghosn E, Lee S, Roan NR. SARS-CoV-2-specific T cells exhibit unique features characterized by robust helper function, lack of terminal differentiation, and high proliferative potential. bioRxiv. 2020:2020.06.08.138826. doi: 10.1101/2020.06.08.138826.

22. Cohen J. COVID-19 shot protects monkeys. Science. 2020;368(6490):456-7. Epub 2020/05/02. doi: 10.1126/science.368.6490.456. PubMed PMID: 32355008.

23. de Wit E, Feldmann F, Cronin J, Jordan R, Okumura A, Thomas T, Scott D, Cihlar T, Feldmann H. Prophylactic and therapeutic remdesivir (GS-5734) treatment in the rhesus macaque model of MERS-CoV infection. Proc Natl Acad Sci U S A. 2020;117(12):6771-6. Epub 2020/02/15. doi: 10.1073/pnas.1922083117. PubMed PMID: 32054787; PMCID: PMC7104368.

24. Bao L, Deng W, Gao H, Xiao C, Liu J, Xue J, Lv Q, Liu J, Yu P, Xu Y, Qi F, Qu Y, Li F, Xiang Z, Yu H, Gong S, Liu M, Wang G, Wang S, Song Z, Zhao W, Han Y, Zhao L, Liu X, Wei Q, Qin C. Reinfection could not occur in SARS-CoV-2 infected rhesus macaques. bioRxiv. 2020:2020.03.13.990226. doi: 10.1101/2020.03.13.990226.

25. Munster VJ, Feldmann F, Williamson BN, van Doremalen N, Perez-Perez L, Schulz J, Meade-White K, Okumura A, Callison J, Brumbaugh B, Avanzato VA, Rosenke R, Hanley PW, Saturday G, Scott D, Fischer ER, de Wit E. Respiratory disease in rhesus macaques inoculated with SARS-CoV-2. Nature. 2020. Epub 2020/05/13. doi: 10.1038/s41586-020-2324-7. PubMed PMID: 32396922.

26. Singh DK, Ganatra SR, Singh B, Cole J, Alfson KJ, Clemmons E, Gazi M, Gonzalez O, Escobedo R, Lee T-H, Chatterjee A, Goez-Gazi Y, Sharan R, Thippeshappa R, Gough M, Alvarez C, Blakley A, Ferdin J, Bartley C, Staples H, Parodi L, Callery J, Mannino A, Klaffke B, Escareno P, Platt RN, Hodara V, Scordo J, Oyejide A, Ajithdoss DK, Copin R, Baum A, Kyratsous C, Alvarez X, Rosas B, Ahmed M, Goodroe A, Dutton J, Hall-Ursone S, Frost PA, Voges AK, Ross CN, Sayers K, Chen C, Hallam C, Khader SA, Mitreva M, Anderson TJC, Martinez-Sobrido L, Patterson JL, Turner J, Torrelles JB, Dick EJ, Brasky K, Schlesinger LS, Giavedoni LD, Carrion R, Kaushal D. SARS-CoV-2 infection leads to acute infection with dynamic cellular and inflammatory flux in the lung that varies across nonhuman primate species. bioRxiv. 2020:2020.06.05.136481. doi: 10.1101/2020.06.05.136481.

27. Chandrashekar A, Liu J, Martinot AJ, McMahan K, Mercado NB, Peter L, Tostanoski LH, Yu J, Maliga Z, Nekorchuk M, Busman-Sahay K, Terry M, Wrijil LM, Ducat S, Martinez DR, Atyeo C, Fischinger S, Burke JS, Slein MD, Pessaint L, Van Ry A, Greenhouse J, Taylor T, Blade K, Cook A, Finneyfrock B, Brown R, Teow E, Velasco J, Zahn R, Wegmann F, Abbink P, Bondzie EA, Dagotto G, Gebre MS, He X, Jacob-Dolan C, Kordana N, Li Z, Lifton MA, Mahrokhian SH, Maxfield LF, Nityanandam R, Nkolola JP, Schmidt AG, Miller AD, Baric RS, Alter G, Sorger PK, Estes JD, Andersen H, Lewis MG, Barouch DH. SARS-CoV-2 infection protects against rechallenge in rhesus macaques. Science. 2020. Epub 2020/05/22. doi: 10.1126/science.abc4776. PubMed PMID: 32434946; PMCID: PMC7243369.

28. Tew JG, Wu J, Fakher M, Szakal AK, Qin D. Follicular dendritic cells: beyond the necessity of T-cell help. Trends Immunol. 2001;22(7):361-7. Epub 2001/06/29. doi: 10.1016/s1471-4906(01)01942-1. PubMed PMID: 11429319.

29. Krautler NJ, Kana V, Kranich J, Tian Y, Perera D, Lemm D, Schwarz P, Armulik A, Browning JL, Tallquist M, Buch T, Oliveira-Martins JB, Zhu C, Hermann M, Wagner U, Brink R, Heikenwalder M, Aguzzi A. Follicular dendritic cells emerge from ubiquitous perivascular precursors. Cell. 2012;150(1):194-206. Epub 2012/07/10. doi: 10.1016/j.cell.2012.05.032. PubMed PMID: 22770220; PMCID: PMC3704230.

30. Long QX, Liu BZ, Deng HJ, Wu GC, Deng K, Chen YK, Liao P, Qiu JF, Lin Y, Cai XF, Wang DQ, Hu Y, Ren JH, Tang N, Xu YY, Yu LH, Mo Z, Gong F, Zhang XL, Tian WG, Hu L, Zhang XX, Xiang JL D. HX, Liu HW, Lang CH, Luo XH, Wu SB, Cui XP, Zhou Z, Zhu MM, Wang J, Xue CJ, Li XF, Wang L, Li ZJ, Wang K, Niu CC, Yang QJ, Tang XJ, Zhang Y, Liu XM, Li JJ, Zhang DC, Zhang F, Liu P, Yuan J, Li Q, Hu JL, Chen J, Huang AL. Antibody responses to SARS-CoV-2 in patients with COVID-19. Nat Med. 2020;26(6):845-8. Epub 2020/05/01. doi: 10.1038/s41591-020-0897-1. PubMed PMID: 32350462.

31. Premkumar L, Segovia-Chumbez B, Jadi R, Martinez DR, Raut R, Markmann A, Cornaby C, Bartelt L, Weiss S, Park Y, Edwards CE, Weimer E, Scherer EM, Rouphael N, Edupuganti S, Weiskopf D, Tse LV, Hou YJ, Margolis D, Sette A, Collins MH, Schmitz J, Baric RS, de Silva AM. The receptor binding domain of the viral spike protein is an immunodominant and highly specific target of antibodies in SARS-CoV-2 patients. Sci Immunol. 2020;5(48). Epub 2020/06/13. doi: 10.1126/sciimmunol.abc8413. PubMed PMID: 32527802.

32. Suthar MS, Zimmerman M, Kauffman R, Mantus G, Linderman S, Vanderheiden A, Nyhoff L, Davis C, Adekunle S, Affer M, Sherman M, Reynolds S, Verkerke H, Alter DN, Guarner J, Bryksin J, Horwath M, Arthur C, Saakadze N, Smith GH, Edupuganti S, Scherer EM, Hellmeister K, Cheng A, Morales JA, Neish AS, Stowell SR, Frank F, Ortlund E, Anderson E, Menachery V, Rouphael N, Metha A, Stephens DS, Ahmed R, Roback J, Wrammert J. Rapid generation of neutralizing antibody responses in COVID-19 patients. medRxiv. 2020:2020.05.03.20084442. doi: 10.1101/2020.05.03.20084442.

33. Liu W, Liu L, Kou G, Zheng Y, Ding Y, Ni W, Wang Q, Tan L, Wu W, Tang S, Xiong Z, Zheng S. Evaluation of Nucleocapsid and Spike Protein-Based Enzyme-Linked Immunosorbent Assays for Detecting Antibodies against SARS-CoV-2. J Clin Microbiol. 2020;58(6). Epub 2020/04/02. doi: 10.1128/JCM.00461-20. PubMed PMID: 32229605; PMCID: PMC7269413.

34. Burbelo PD, Riedo FX, Morishima C, Rawlings S, Smith D, Das S, Strich JR, Chertow DS, Davey RT, Cohen JI. Detection of Nucleocapsid Antibody to SARS-CoV-2 is More Sensitive than Antibody to Spike Protein in COVID-19 Patients. medRxiv. 2020:2020.04.20.20071423. doi: 10.1101/2020.04.20.20071423.

35. Vardhana SA, Wolchok JD. The many faces of the anti-COVID immune response. J Exp Med. 2020;217(6). Epub 2020/05/01. doi: 10.1084/jem.20200678. PubMed PMID: 32353870.

36. Geissmann F, Jung S, Littman DR. Blood monocytes consist of two principal subsets with distinct migratory properties. Immunity. 2003;19(1):71-82. Epub 2003/07/23. doi: 10.1016/s1074-7613(03)00174-2. PubMed PMID: 12871640.

37. Patel AA, Zhang Y, Fullerton JN, Boelen L, Rongvaux A, Maini AA, Bigley V, Flavell RA, Gilroy DW, Asquith B, Macallan D, Yona S. The fate and lifespan of human monocyte subsets in steady state and systemic inflammation. J Exp Med. 2017;214(7):1913-23. Epub 2017/06/14. doi: 10.1084/jem.20170355. PubMed PMID: 28606987; PMCID: PMC5502436.

38. Yang J, Zhang L, Yu C, Yang XF, Wang H. Monocyte and macrophage differentiation: circulation inflammatory monocyte as biomarker for inflammatory diseases. Biomark Res. 2014;2(1):1. Epub 2014/01/09. doi: 10.1186/2050-7771-2-1. PubMed PMID: 24398220; PMCID: PMC3892095.

39. Cervantes-Barragan L, Zust R, Weber F, Spiegel M, Lang KS, Akira S, Thiel V, Ludewig B. Control of coronavirus infection through plasmacytoid dendritic-cell-derived type I interferon. Blood. 2007;109(3):1131-7. Epub 2006/09/21. doi: 10.1182/blood-2006-05-023770. PubMed PMID: 16985170.

40. Lv H, Wu NC, Tsang OT, Yuan M, Perera R, Leung WS, So RTY, Chan JMC, Yip GK, Chik TSH, Wang Y, Choi CYC, Lin Y, Ng WW, Zhao J, Poon LLM, Peiris JSM, Wilson IA, Mok CKP. Cross-reactive antibody response between SARS-CoV-2 and SARS-CoV infections. bioRxiv. 2020. Epub 2020/06/09. doi: 10.1101/2020.03.15.993097. PubMed PMID: 32511317; PMCID: PMC7239046.

41. Robbiani DF, Gaebler C, Muecksch F, Lorenzi JCC, Wang Z, Cho A, Agudelo M, Barnes CO, Gazumyan A, Finkin S, Hägglöf T, Oliveira TY, Viant C, Hurley A, Hoffmann H-H, Millard KG, Kost RG, Cipolla M, Gordon K, Bianchini F, Chen ST, Ramos V, Patel R, Dizon J, Shimeliovich I, Mendoza P, Hartweger H, Nogueira L, Pack M, Horowitz J, Schmidt F, Weisblum Y, Michailidis E, Ashbrook AW, Waltari E, Pak JE, Huey-Tubman KE, Koranda N, Hoffman PR, West AP, Rice CM, Hatziioannou T, Bjorkman PJ, Bieniasz PD, Caskey M, Nussenzweig MC. Convergent antibody responses to SARS-CoV-2 in convalescent individuals. Nature. 2020. doi: 10.1038/s41586-020-2456-9.

42. Seydoux E, Homad LJ, MacCamy AJ, Parks KR, Hurlburt NK, Jennewein MF, Akins NR, Stuart AB, Wan YH, Feng J, Whaley RE, Singh S, Boeckh M, Cohen KW, McElrath MJ, Englund JA, Chu HY, Pancera M, McGuire AT, Stamatatos L. Analysis of a SARS-CoV-2-Infected Individual Reveals Development of Potent Neutralizing Antibodies with Limited Somatic Mutation. Immunity. 2020. Epub 2020/06/21. doi: 10.1016/j.immuni.2020.06.001. PubMed PMID: 32561270.

43. Salazar E, Perez KK, Ashraf M, Chen J, Castillo B, Christensen PA, Eubank T, Bernard DW, Eagar TN, Long SW, Subedi S, Olsen RJ, Leveque C, Schwartz MR, Dey M, Chavez-East C, Rogers J, Shehabeldin A, Joseph D, Williams G, Thomas K, Masud F, Talley C, Dlouhy KG, Lopez BV, Hampton C, Lavinder J, Gollihar JD, Maranhao AC, Ippolito GC, Saavedra MO, Cantu CC, Yerramilli P, Pruitt L, Musser JM. Treatment of Coronavirus Disease 2019 (COVID-19) Patients with Convalescent Plasma. Am J Pathol. 2020. Epub 2020/05/31. doi: 10.1016/j.ajpath.2020.05.014. PubMed PMID: 32473109; PMCID: PMC7251400.

44. Chen L, Xiong J, Bao L, Shi Y. Convalescent plasma as a potential therapy for COVID-19. Lancet Infect Dis. 2020;20(4):398-400. Epub 2020/03/03. doi: 10.1016/S1473-3099(20)30141-9. PubMed PMID: 32113510.

45. Roback JD, Guarner J. Convalescent Plasma to Treat COVID-19: Possibilities and Challenges. JAMA. 2020. Epub 2020/03/29. doi: 10.1001/jama.2020.4940. PubMed PMID: 32219429.

46. Gharbharan A, Jordans CCE, GeurtsvanKessel C, den Hollander JG, Karim F, Mollema FPN, Stalenhoef JE, Dofferhoff A, Ludwig I, Koster A, Hassing R-J, Bos JC, van Pottelberge GR, Vlasveld IN, Ammerlaan HSM, Segarceanu E, Miedema J, van der Eerden M, Papageorgiou G, te Broekhorst P, Swaneveld FH, Katsikis PD, Mueller Y, Okba NMA, Koopmans MPG, Haagmans BL, Rokx C, Rijnders B. Convalescent Plasma for COVID-19. A randomized clinical trial. medRxiv. 2020:2020.07.01.20139857. doi: 10.1101/2020.07.01.20139857.

47. SARS-CoV-2 Surveillance for a Nonhuman Primate Breeding Research Facility. Journal of Medical Primatology. doi: 10.1111/jmp.12483.

48. Sanville B, Corbett R, Pidcock W, Hardin K, Sebat C, Nguyen MV, Thompson GR, Haczku A, Schivo M, Cohen S. A Community Transmitted Case of Severe Acute Respiratory Distress Syndrome due to SARS CoV2 in the United States. Clin Infect Dis. 2020. Epub 2020/04/01. doi: 10.1093/cid/ciaa347. PubMed PMID: 32227197; PMCID: PMC7197621.

49. Verma A, Schmidt BA, Elizaldi SR, Nguyen NK, Walter KA, Beck Z, Trinh HV, Dinasarapu AR, Lakshmanappa YS, Rane NN, Matyas GR, Rao M, Shen X, Tomaras GD, LaBranche CC, Reimann KA, Foehl DH, Gach JS, Forthal DN, Kozlowski PA, Amara RR, Iyer SS. Impact of Th1 CD4 Follicular Helper T Cell Skewing on Antibody Responses to an HIV-1 Vaccine in Rhesus Macaques. J Virol. 2020;94(6). Epub 2019/12/13. doi: 10.1128/JVI.01737-19. PubMed PMID: 31827000; PMCID: PMC7158739.

50. Havenar-Daughton C, Reiss SM, Carnathan DG, Wu JE, Kendric K, Torrents de la Pena A, Kasturi SP, Dan JM, Bothwell M, Sanders RW, Pulendran B, Silvestri G, Crotty S. Cytokine-Independent Detection of Antigen-Specific Germinal Center T Follicular Helper Cells in Immunized Nonhuman Primates Using a Live Cell Activation-Induced Marker Technique. J Immunol. 2016;197(3):994-1002. Epub 2016/06/24. doi: 10.4049/jimmunol.1600320. PubMed PMID: 27335502; PMCID: PMC4955744.

51. Roederer M, Nozzi JL, Nason MC. SPICE: exploration and analysis of post-cytometric complex multivariate datasets. Cytometry A. 2011;79(2):167-74. Epub 2011/01/26. doi: 10.1002/cyto.a.21015. PubMed PMID: 21265010; PMCID: PMC3072288.

52. Kozlowski PA, Lynch RM, Patterson RR, Cu-Uvin S, Flanigan TP, Neutra MR. Modified wick method using Weck-Cel sponges for collection of human rectal secretions and analysis of mucosal HIV antibody. J Acquir Immune Defic Syndr. 2000;24(4):297-309. Epub 2000/10/03. doi: 10.1097/00126334-200008010-00001. PubMed PMID: 11015145.

53. Ng D, Goldgof G, Shy B, Levine A, Balcerek J, Bapat SP, Prostko J, Rodgers M, Coller K, Pearce S, Franz S, Du L, Stone M, Pillai S, Sotomayor-Gonzalez A, Servellita V, Sanchez-San Martin C, Granados A, Glasner DR, Han LM, Truong K, Akagi N, Nguyen DN, Neumann N, Qazi D, Hsu E, Gu W, Santos YA, Custer B, Green V, Williamson P, Hills NK, Lu CM, Whitman JD, Stramer S, Wang C, Reyes K, Hakim J, Sujishi K, Alazzeh F, Pharm L, Oon C-Y, Miller S, Kurtz T, Hackett J, Simmons G, Busch MP, Chiu CY. SARS-CoV-2 seroprevalence and neutralizing activity in donor and patient blood from the San Francisco Bay Area. medRxiv. 2020:2020.05.19.20107482. doi: 10.1101/2020.05.19.20107482.

