## Supplementary table and figure legends for "SARS-CoV-2 infection induces germinal center responses with robust stimulation of CD4 T follicular helper cells in rhesus macaques"

**SUPPLEMENTARY FIGURE TITLES AND LEGENDS**

**Key Resources Table. Antibody and chemical information.** All antibodies and chemicals used in the study are denoted.

**Supplementary Table 1. Animal information.** Sex, Age, Body weight, treatment, infusion volume, blood donor type, total volume infused, necropsy date, prior treatments and extra notes for all animals on the study.

**Supplementary Table 2. Spleen AIM Assay conditions.** N, S, M, ORF (nsp3, nsp4, 3a, 8), or P/I stimulation conditions for each animal on the study.

**Supplementary Table 3. Mediastinal LN AIM Assay conditions.** N or S peptide stimulation conditions for mediastinal LN for each animal on the study.

**Supplemental Figure 1. Clinical symptoms and complete blood counts following SARS-CoV-2 infection (A)** Alveolar septae are expanded by a mixed inflammatory cell infiltrate and alveoli contain macrophages [black arrow] and occasional neutrophils; interstitial thickening is also apparent [blue arrow]. **(B)** Rectal Temperature [°F] and Body weight [kg] of SARS-CoV-2 infected rhesus macaques over the course of the study **(C)** Frequency of pro-inflammatory monocytes [CD14+CD16+] expressing CX3CR1 at day 0 and 2 within blood following infection **(D)** RBC Counts [x10^6 /ul blood], Platelets [x10^5 /ul blood], WBC counts [x10^3 /ul blood], Lymphocyte counts [/ul blood] and Monocyte counts [/ul blood] over the course of the study; Infected (black circles) are RM that were infected and received no plasma treatment, I+CP (blue circles) are RM that were infected and received human convalescent plasma, and I+NP are RM that were infected and received normal plasma from patients with no history of SARS-CoV-2 infection.

**Supplemental Figure 2. SARS-CoV-2 infection elicits Ki67+PD-1+ Th1 cells at Day 7 that are polyfunctional** **(A)** tSNE plots of CD4+Ki-67+PD-1+ CXCR5+/CXCR5- populations show representation of Th1 effectors [CXCR3^+^], Th2 effectors [CCR4^+^], and Th17 effectors [CCR6^+^] **(B)** Increase of Ki67^+^PD-1^+^ Th1 cells in both cell frequency and cells/ul blood at day 7 post infection **(C)** Kinetics of Ki-67^+^ PD-1^+^ Th2 cells [/ul blood] **(D)** Kinetics of Ki-67^+^ PD-1^+^ Th17 cells [/ul blood] **(E)** Representative expression of IFNg, IL-17, IL-2, TNF-α within two CD4 populations: CD107a/b^+^ cells and IL-21^+^ cells after stimulation **(F)** Serum CXCL13 following infection

**Supplemental Figure 3.** **Splenic GC TFH, Tfr, and Treg populations during SARS-CoV-2 infection** **(A)** Flow Plot of splenic GC TFH cells [CXCR5^+^PD-1^hi^] at necropsy (d11-d14pi) and dot plot graph designating an increase of GC TFH in SARS-CoV-2 infected rhesus macaques **(B)** Flow Plot of Treg [Foxp3^+^] and Tfr cells. Controls (circles) are RM that were infected and received no plasma treatment, Infused (triangles) are RM that were infected and received either convalescent plasma or normal plasma.

**Supplemental Figure 4. SARS-CoV-2 infection increases mediastinal lymph node size and elicits GC Tfh cells. (A)** Large aggregate of bronchial associated lymphoid tissue [arrow] adjacent to a moderately large airway and associated blood vessel. Two well defined germinal centers can be seen within the BALT **(B)** Comparison of SARS-CoV-2 uninfected [left] and SARS-CoV-2 infected [right] Mediastinal lymph nodes show distinct lymphadenopathy **(C)** Detection of follicular dendritic cells (FDCs) gating on CD45^-^Lin^-^CD14^-^CD35^+^CD21^+^ using flow cytometry; three different clones are used for comparison [BL13, BU32, B-LY4] **(D)** Correlations between % total FDCs and % total GC T_FH_ or % total GC B cells **(E)** Peptide pool stimulation if mediastinal lymph node cells with Spike (S), Nucleocapsid (N), Membrane protein (M), or PMA/Iono (P/I) show responses in CXCR5^+^, CXCR5^-^, or Naïve CD4 T cell populations

**KEY RESOURCES TABLE**

1. **ANTIBODIES**

| **Reagents** | **Clone** | **Source** | **Identifier (Cat#)** |
| --- | --- | --- | --- |
| CD3-AF 700 | SP34-2 | BD Biosciences | 557917 |
| CD3-APC-Cy7 | SP34-2 | BD Biosciences | 557757 |
| CD4-BV 650 | L200 | BD Biosciences | 563737 |
| CD8-BV 510 | SK-1 | BD Biosciences | 563919 |
| CD8-BUV 805 | SK-1 | BD Biosciences | 564913 |
| CD20-APC-Cy7 | 2H7 | BioLegend | 302314 |
| CD20-BV 421 | 2H7 | BioLegend | 302328 |
| CD95-BUV 737 | DX2 | BD Biosciences | 564710 |
| CD279(PD-1)-Pe-Cy7 | EH12.2H8 | BioLegend | 329918 |
| CX3CR1- PE-CF594 | 2A9-1 | BioLegend | 341624 |
| CXCR3-BV 786 | IC6 | BD Biosciences | 741005 |
| CXCR5-PE | MU5UBEE | Thermofischer | 12-9185-411G1 |
| CCR4-BV 605 | 1G1 | BD Biosciences | 562906 |
| CCR6-PE-CF594/A610 | G034E3 | BioLegend | 353430 |
| CCR7-BV 650 | 3D12 | BD Biosciences | 563407 |
| HLA-DR-BV 786 | L243 | BioLegend | 307642 |
| CD69-BV 711 | FN50 | BioLegend | 310944 |
| CD69-Pe-Cy7 | FN50 | Invitrogen | 25-0699-42 |
| CD14-AF 700 | MSE2 | BD Biosciences | 301822 |
| CD16-BV 605 | 3G8 | BD Biosciences | 563172 |
| CD11b-BV 510 | ICRF44 | Thermofisher | 563088 |
| CD11c-Pe-Cy7 | 3.9 | Invitrogen | 25-0116-42 |
| CD103-APC | 2G5 | Beckman Coulter | B06204 |
| CD66-APC | TET2 | Miltenyi | 130-118-539 |
| CD163-PE | GHI/61 | BioLegend | 333606 |
| CD123-BV 421 | 7G3 | Thermofisher | 501129764 |
| Granzyme B-BV 421 | GB11 | BioLegend | 515408 |
| CD80-AF 488 | 2D10.4 | Invitrogen | 11-0809-42 |
| CD86-AF 488 | IT2.2 | BioLegend | 305414 |
| Ki-67-AF 488 | B56 | BD Biosciences | 558616 |
| Ki-67-BV 510 | B56 | BD Biosciences | 563462 |
| CD28-Pe-Cy7 | CD28.2 | Tonbo | 40-0289-U500 |
| a4b7-PE | Act-1 | NHP Reag Res | PR-1422 |
| CD45-AF 488 | D058-1283 | BD Biosciences | 557803 |
| CD45-BV 605 | D058-1283 | BD Biosciences | 564098 |
| CD140b-APC | 18A2 | BioLegend | 323608 |
| Bcl-6-APC-Cy7 | K112-91 | BD Biosciences | 563581 |
| CD21- PE-CF594 | B-ly4 | BD Biosciences | 563474 |
| SLAM-AF 488 | A12(7D4) | BioLegend | 306312 |
| Foxp3-APC | 206D | BioLegend | 320114 |
| CD278 (ICOS)-BV 785 | C396.4A | BioLegend | 313534 |
| CD25-APC | BC96 | Tonbo | 20-0259-T100 |
| CD 134 (OX40)-BV 786 | L106 | BD Biosciences | 744746 |
| 4-1BB-AF 488 | 4B4-1 | BD Biosciences | 11-1379-42 |
| TNFa-AF 488 | Mab11 | BioLegend | 502906 |
| IL-21-APC | 3A3-N2.1 | BD Biosciences | 560493 |
| IFNG-PeCy7 | B27 | BioLegend | 506518 |
| IL2- PE-CF594 | MO1-17H12 | BioLegend | 500344 |
| CD107a-PE | H4A3 | BioLegend | 328608 |
| CD107b-PE | EbioH4B4 | Thermofischer | 12-1078-42 |
| IL-17-BV 421 | eBio64DEC17 | eBiosciences | 48-7179-42 |
| CD45 RO- PE-CF594 | UCHL-1 | BD Biosciences | 562299 |
| CD28-Purified Ab | CD28.2 | BD Biosciences | 555725 |
| CD49d-Purified Ab | 9F10 | BD Biosciences | 555501 |
| Live/dead-APC-Cy7 |  | Life technologies | L34976 |
| Live/dead-BV 510 |  | Life technologies | L34966 |

2. **Chemicals:**

| **Reagents** | **Source** | **Identifier (Cat#)** | **Concentrations** |
| --- | --- | --- | --- |
| Golgi Stop | BD Biosciences | 554724 |  |
| Golgi Plug | BD Biosciences | 555029 |  |
| PMA+ Ionomycin cocktail mix | eBiosciences | 00-4970-93 | 500X |
| T cell activation/ expansion kit NHP | MACS Miltenyi Biotech | 130-092-919 |  |
| AIM V media | Gibco | 12055091 |  |
| RPMI media-1640 | Gibco |  |  |
| Cytofix/cytoperm | BD Biosciences | 51-2090K2 |  |
| DNAse-I | Roche diagnostics | BM070 |  |
| Collagenase-type IV | Worthington Biomedical corporation | LS004188 |  |
