## Supplementary material for "SARS-CoV-2 infection induces germinal center responses with robust stimulation of CD4 T follicular helper cells in rhesus macaques": Animal Information

| Supplementary Table 1. Animal information | | | | | | | | | |
|---|---|---|---|---|---|---|---|---|---|
| **Animal code** | **Sex** | **Age** | **Body Weight (kg)** | **Treatment** | **Infusion Volume** | **Total volume infused (4ml/kg)** | **Nx Day** | **Prior treatments** | Clinical Notes |
| Control 1 | **F** | 5:10:09 | **6** | **no plasma** | **N/A** | **N/A** | D11 | 44470 received clinical analgesics and antibiotics for trauma and nutritional supplements due to lean BCS | |
| Control 2 | **F** | 4:10:07 | **7.01** | **no plasma** | **N/A** | **N/A** | D11 | | Sneezed during study |
| CP1 | **M** | 5:10:14 | **8.43** | **Conval. Plasma** | **27ml** | **33.7** | D12 | Dexamethasone suppression/ACTH stimulation tests as part of BBA testing (2014) Experimental vaccine (for Campylobacter coli in 2018) | |
| NP1 | **M** | 5:10:26 | **8.95** | **normal plasma** | **27ml** | **35.8** | D12 | 44309 has historically received clinical analgesics, antibiotics, and supplements for trauma cases | |
| NP2 | **M** | 5:09:18 | **9.74** | **normal plasma** | **30ml** | **39.0** | D13 | | |
| CP2 | **M** | 5:10:19 | **8.83** | **Conval. Plasma** | **27ml** | **35.3** | D13 | 44379 has received analgesics/antibiotics for trauma and antibiotics and probiotics for diarrhea historically | Sneezed during study |
| Control 3 | **F** | 4:10:23 | **5.47** | **no plasma** | **N/A** | **N/A** | D14 | 45159 has received analgesics/antibiotics for trauma and probiotics for diarrhea | Dermatitis |
| Control 4 | **M** | 4:10:00 | **10.72** | **no plasma** | **N/A** | **N/A** | D14 | 45359 has received analgesics/antibiotics for trauma and probiotics for diarrhea | |
