## Supplementary material for "SARS-CoV-2 infection induces germinal center responses with robust stimulation of CD4 T follicular helper cells in rhesus macaques": Spleen AIM Assay conditions

| Supplementary Table 2. Spleen AIM Assay conditions | | | | | | | | | |
|---|---|---|---|---|---|---|---|---|---|
| **Spleen** | **Animal code** | **N** | **S** | **M** | **ORF1-nsp3** | **ORF1-nsp4** | **ORF3a** | **ORF8** | **P/I** |
| **1** | Control 1 | Excluded due to low CD3 events | √ | √ | √ | √ | √ | √ | √ |
| **2** | Control 2 | √ | √ | √ | √ | √ | √ | √ | √ |
| **3** | CP1 | √ | √ | √ | √ | √ | √ | √ | √ |
| **4** | NP1 | √ | √ | √ | √ | √ | √ | √ | √ |
| **5** | NP2 | Excluded due to low recovery of CD95+ cells | | | | | | | |
| **6** | CP2 | | | | | | | | |
| **7** | Control 3 | √ | √ | √ | √ | √ | √ | √ | √ |
| **8** | Control 4 | √ | √ | √ | √ | √ | √ | √ | √ |
