## Supplementary material for "SARS-CoV-2 infection induces germinal center responses with robust stimulation of CD4 T follicular helper cells in rhesus macaques": Mediastinal LN AIM Assay conditions

| Supplementary Table 3. Mediastinal LN AIM Assay conditions | | | |
|---|---|---|---|
| **Med LN** | **Animal code** | **N** | **S** |
| **1** | Control 1 | Excluded due to low recovery of CD95+ cells | |
| **2** | Control 2 | | |
| **3** | CP1 | ✓ | ✓ |
| **4** | NP1 | ✓ | ✓ |
| **5** | NP2 | Excluded due to low recovery of CD95+ cells | |
| **6** | CP2 | | |
| **7** | Control 3 | ✓ | ✓ |
| **8** | Control 4 | ✓ | ✓ |
