## Supplementary figures and images for "SARS-CoV-2 infection induces germinal center responses with robust stimulation of CD4 T follicular helper cells in rhesus macaques"

### Clinical symptoms and complete blood counts following SARS-CoV-2 infection

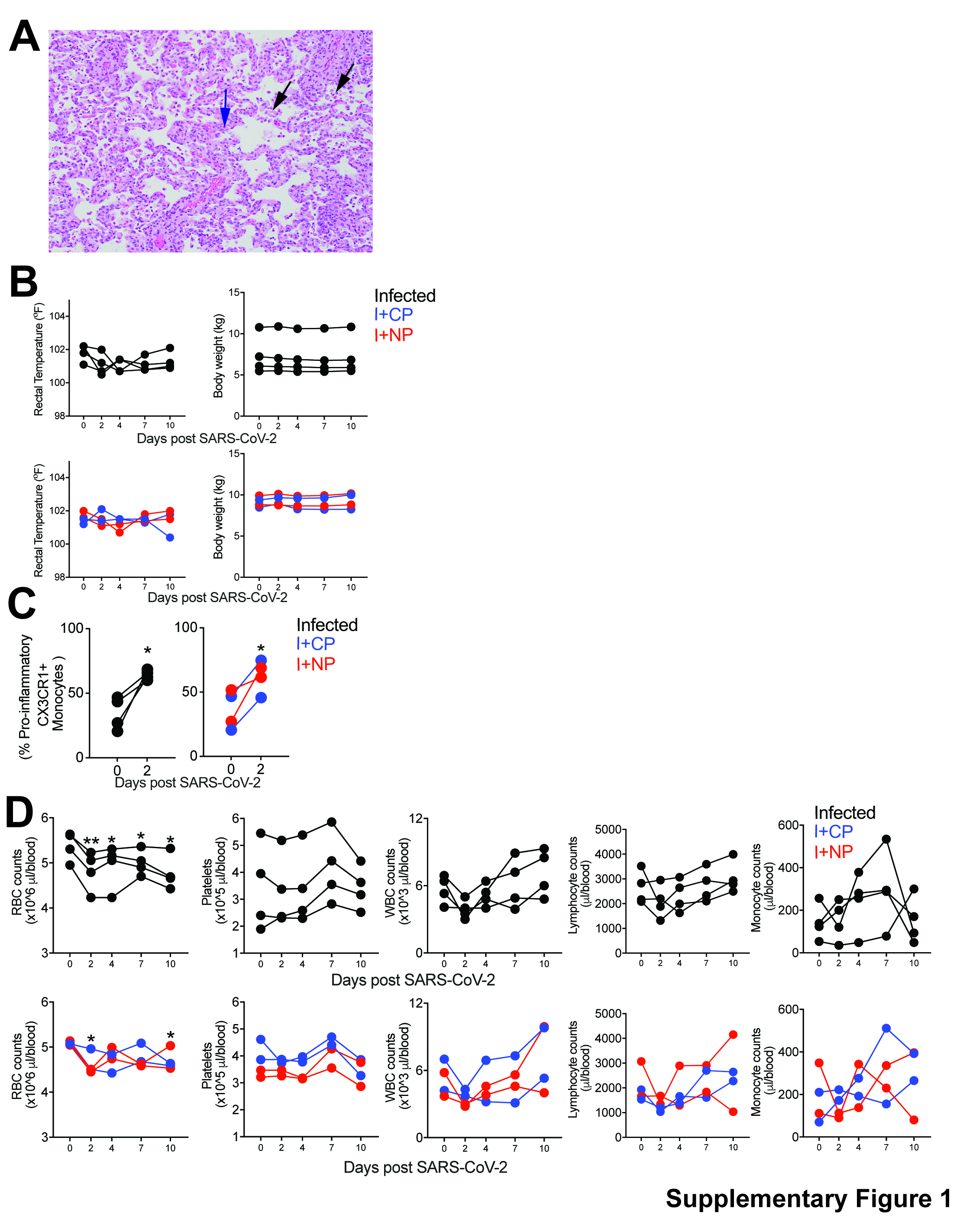

### SARS-CoV-2 infection elicits Ki67+PD-1+ Th1 cells at Day 7 that are polyfunctional

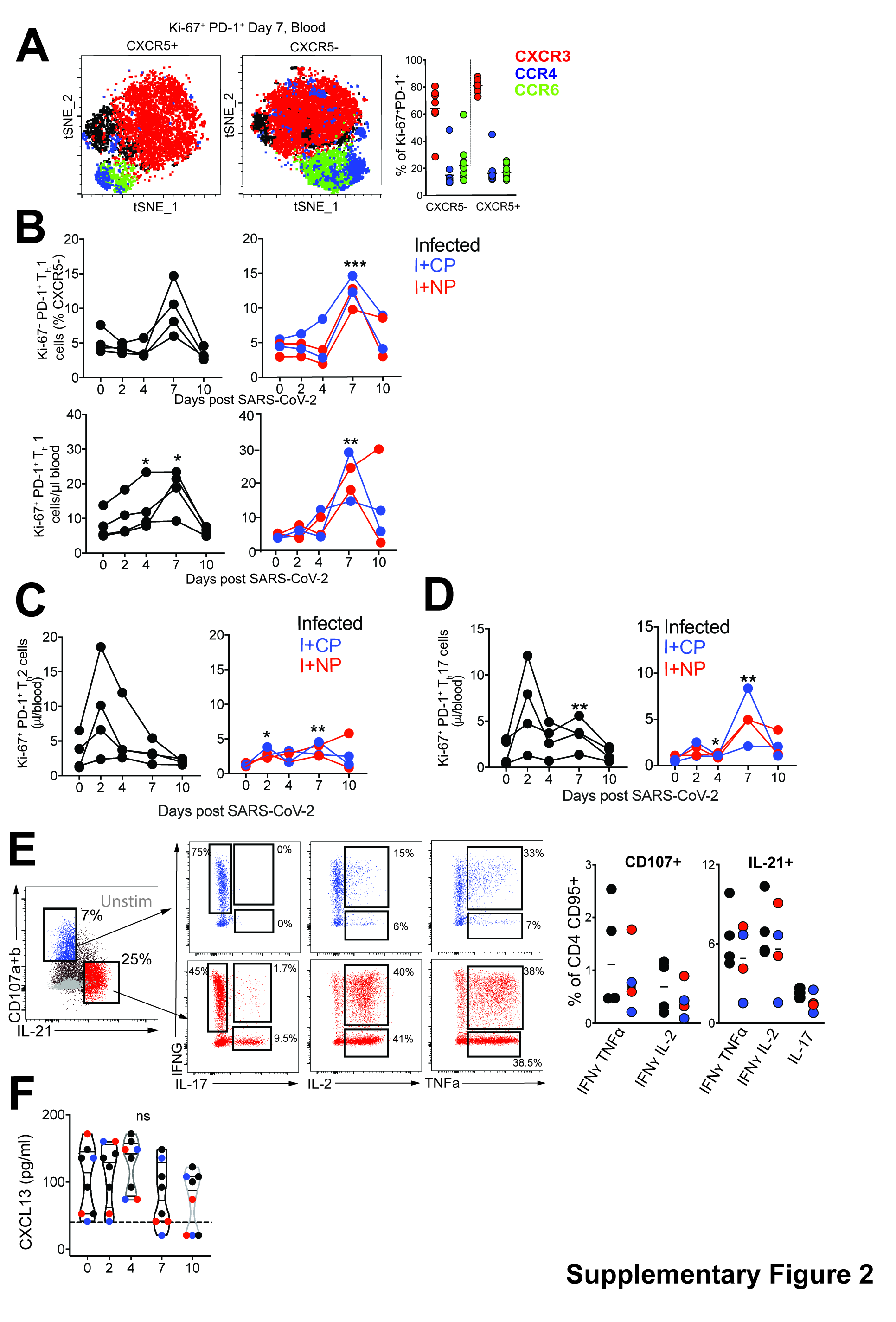

### SARS-CoV-2 infection increases mediastinal lymph node size and peptide pool stimulation affect on CXCR5+, CXCR5- and Naive T cells.

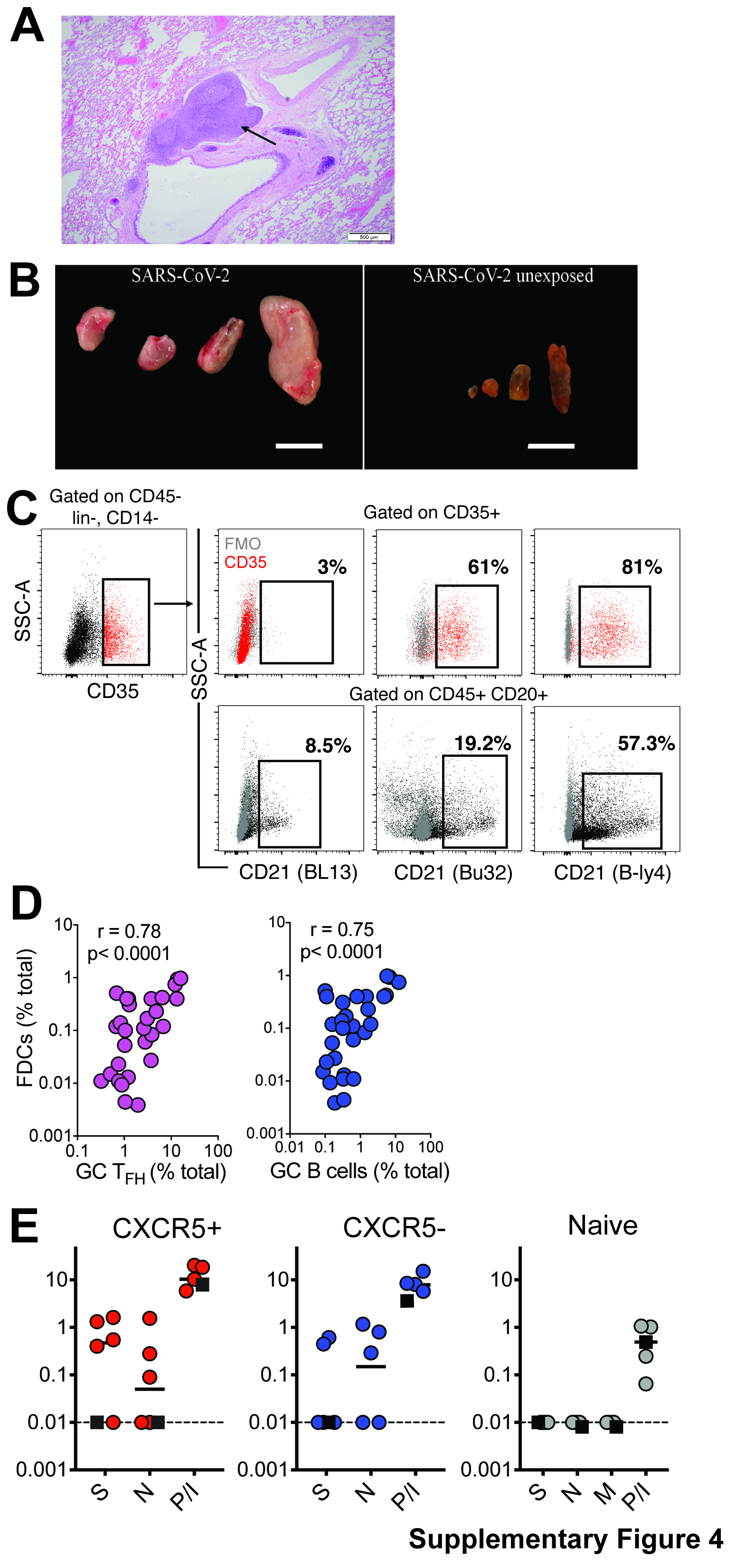

### Splenic GC TFH, Tfr, and Treg populations during SARS-CoV-2 infection

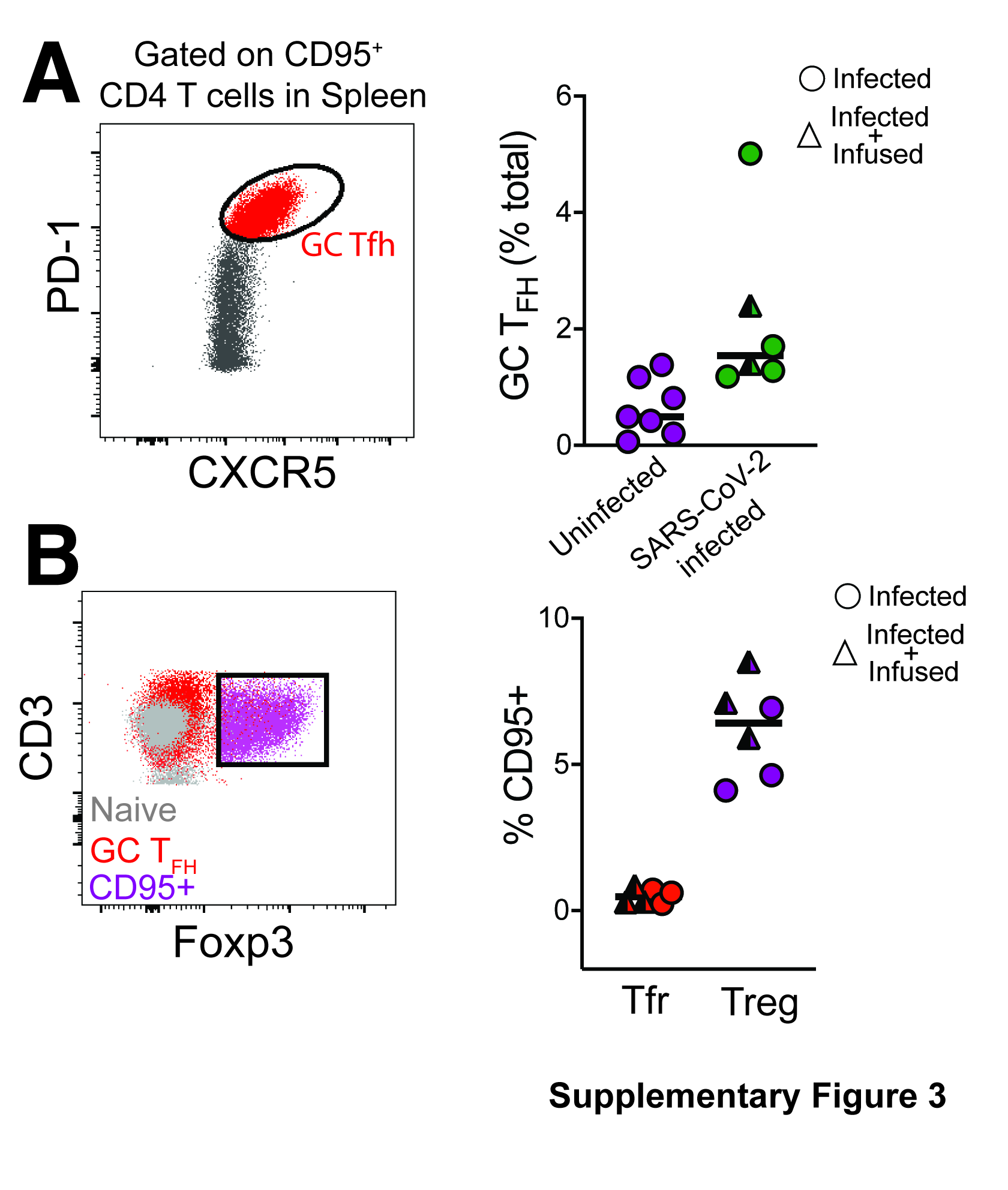
